# Cup is essential for *oskar* mRNA translational repression during early *Drosophila* oogenesis

**DOI:** 10.1101/2023.01.27.525950

**Authors:** Livia V. Bayer, Samantha Milano, Stephen K. Formel, Harpreet Kaur, Rishi Ravichandran, Juan A. Cambeiro, Lizaveta Slinko, Irina E. Catrina, Diana P. Bratu

## Abstract

The proper timing of mRNA translation is crucial across many biological systems for processes such as intercellular communication, body pattern formation, and morphogenesis. The main *D. melanogaster* posterior determinant, *oskar*, is maternally transcribed, but only translated when properly localized at the oocyte’s posterior cortex. Bruno 1 and Cup are two effector proteins known to participate in multiple aspects of *oskar* mRNA regulation. Current model describes a mechanism in which Bruno 1 is necessary for Cup’s recruitment to *oskar* mRNA, and Bruno 1 is indispensable for its translational repression. Here, we reveal that the Bruno 1-Cup interaction, as well as their interdependent influence on each other’s mRNA and protein expression, lead to precise *oskar* mRNA regulation during early oogenesis. We show that these factors stably associate with the *oskar* mRNA *in vivo*, but surprisingly, Bruno 1’s stable association with *oskar* mRNA depends on Cup, while Bruno 1 is not necessary for Cup association to *oskar* mRNA. During early oogenesis, Cup, not Bruno 1, is the essential factor for *oskar* mRNA repression. Cup is a crucial P-body member that maintains proper P-body morphology during oogenesis, as well as it is necessary for *oskar* mRNA’s association with P-bodies, thus driving the translational repression and stability of *oskar* mRNA. Our experimental results collectively suggest a regulatory mechanism where a feedback loop between Bruno 1 and Cup coordinates *oskar* mRNA regulation in the egg chamber allowing for proper development to occur.

## INTRODUCTION

Post-transcriptional gene regulation is important for complex processes, such as cell motility, synaptic plasticity, and organism development. This regulation is primarily dictated by the dynamic associations between messenger RNA (mRNA) and proteins (mRNP) throughout the mRNA’s life. In the *Drosophila melanogaster* egg chamber, *oskar (osk)* is critical for establishing the developmental patterning along with the future germline (Lehmann and Nusslein-Volhard 1986). A myriad of proteins form a large mRNP involved in the stability, transport, localization, translational repression, and translational activation of *osk* mRNA. Two of these proteins, Cup and Bruno 1 (Bru1), have been proven necessary for precise *osk* mRNA regulation (Kim et al. 2015; Kim-Ha, Kerr, and Macdonald 1995; Akira Nakamura, Sato, and Hanyu-Nakamura 2004; Wilhelm et al. 2003). However, the dynamics of their spatial and temporal interactions have remained elusive. We now show that both Bru1 and Cup associate with *osk* mRNP in the nurse cells and travel together into the oocyte.

Bru1 is a member of the CELF superfamily (CUGBP and Elav-like family) which directly binds *osk* mRNA’s 3’ UTR at two regions: a combined A|B site and a C site, where multiple Bru1 response elements (BREs) are located (Kim-Ha, Kerr, and Macdonald 1995; M. Snee et al. 2008). Repression of *osk* mRNA translation requires both of these regions, and the importance of BREs was demonstrated by several studies carried out during mid-to-late oogenesis (Kim-Ha, Kerr, and Macdonald 1995; Reveal et al. 2010). Interestingly, the C site is also involved in the translational activation of *osk* mRNA at the posterior of the oocyte (Reveal et al. 2010), yet whether this activation is dependent on *osk’*s localization is not fully understood. Notably, it has also been shown that Bru1 is able to repress in a BRE independent manner, *germ cell-less* mRNA, starting only at stage 4/5 of oogenesis (Moore, Han, and Lasko 2009).

Bru1 has two RNA-recognition domains, RRM 1+2, and an extended RRM 3 that are important for binding *osk* mRNA and other maternal mRNAs *in vitro* (Reveal et al. 2011). Bru1 can also dimerize through its N-terminal domain and possibly forms large silencing particles leading to *osk* mRNA translational repression (Chekulaeva, Hentze, and Ephrussi 2006; Kim et al. 2015). Without this domain, *osk* mRNA granules do not form, and *osk* mRNA is not translated (Bose et al. 2022). Interestingly, this is also one of the domains where Bru1 directly binds Cup with very high affinity (Kim et al. 2015).In addition, Bru1 expression must be maintained at a critical threshold, as either loss-of-function or overexpression of Bru1 led to similar defects, both being detrimental for egg chamber development (M.J. Snee et al. 2007; M. Snee et al. 2008).

Cup is a canonical eIF4E binding protein (4E-BP) that blocks translation initiation of mRNAs (Wilhelm et al. 2003; Kinkelin et al. 2012; Akira Nakamura, Sato, and Hanyu-Nakamura 2004). Cup also interacts with the CCR4-Not complex to allow for the deadenylation of mRNAs while simultaneously blocking decapping enzymes to maintain a translationally silent and stable mRNA (Igreja and Izaurralde 2011). For *osk* mRNA, the current model suggests that Bru1 recruits Cup to the transcript, forming a Bru1-Cup-eIF4E translational repression complex (Akira Nakamura, Sato, and Hanyu-Nakamura 2004; Wilhelm et al. 2003). We show that Cup is able to join *osk* mRNP and actually repress its translation in the absence of Bru1, and surprisingly, Cup is necessary to stabilize the interaction between *osk* mRNA and Bru1. Interestingly, we reveal a strict codependence between Bru1 and Cup stability and distribution in the egg chamber.

Lastly, Cup has been identified as a processing body (P-body) component due to its association with two core P-body proteins, Me31B and Trailer Hitch (Tral) (Akira Nakamura, Sato, and Hanyu-Nakamura 2004; Wang et al. 2017). Our investigation on the connection between Cup and *osk* mRNA translational repression identified Cup as a crucial cellular component that regulates Me31B-condensates physical state, which in turn, possibly ensures proper P-body function.

P-bodies are membraneless organelles that form via liquid-liquid phase separation. They are composed of proteins that contain intrinsically disordered regions (IDR), mRNAs, and mRNA-binding factors (reviewed in (Standart and Weil 2018)). Formation of P-body condensates is not fully understood, but their integrity and the physical state they adopt are regulated by structurally distinct proteins, such as Me31B and Tral (Sankaranarayanan et al. 2021). Though previously thought to be sites of mRNA turnover, recent research demonstrates that P-bodies are also storage hubs for translationally silenced mRNAs (reviewed in (Luo, Na, and Slavoff 2018)) which get released into the cytoplasm upon developmental cues or as a result of disrupted condensate organization (Sankaranarayanan et al. 2021). *osk* mRNA is one of the several post-transcriptionally regulated mRNAs stored in P-bodies (Nakamura et al. 2001; Weil et al. 2012). Here we bring new insights into the regulation of *osk* mRNA translational repression and association with P-bodies, that has not been addressed before during early oogenesis. Particularly we investigated the role of Bru1 and Cup in *osk* mRNA recruitment into P-bodies during early oogenesis.

## RESULTS

### Bru1 and Cup maintain a stable association with *osk* mRNA in the egg chamber

The *Drosophila melanogaster* egg chamber is an excellent system to study important details of the spatio-temporal events that occur during post-transcriptional regulation. *Drosophila* oogenesis spans from the germarium through 14 stages of egg-chamber development that here we referred to as early (1-4), mid (5-9) and late (10-14) (Fig. 1A). The egg chamber consists of 16 germline cells, including a single oocyte and 15 nurse cells surrounded by somatic follicle cells (reviewed in (McLaughlin and Bratu 2015)). Nurse cells supply organelles, factors, and key transcripts to the transcriptionally repressed oocyte. *osk* mRNA moves as a translationally silenced mRNP from the nurse cells into the oocyte, where it eventually anchors tightly at the posterior cortex (Fig. 1A green). Translation of *osk* mRNA occurs only at the posterior pole (Fig. 1A red). Premature or ectopic expression of Osk is lethal to the embryo (Ephrussi and Lehmann 1992; Kim-Ha J 1991). Moreover, *osk* mRNA itself is crucial for progression through oogenesis, with its absence leading to arrest in egg chamber development during mid-oogenesis (Kanke et al. 2015).

**Figure 1.**
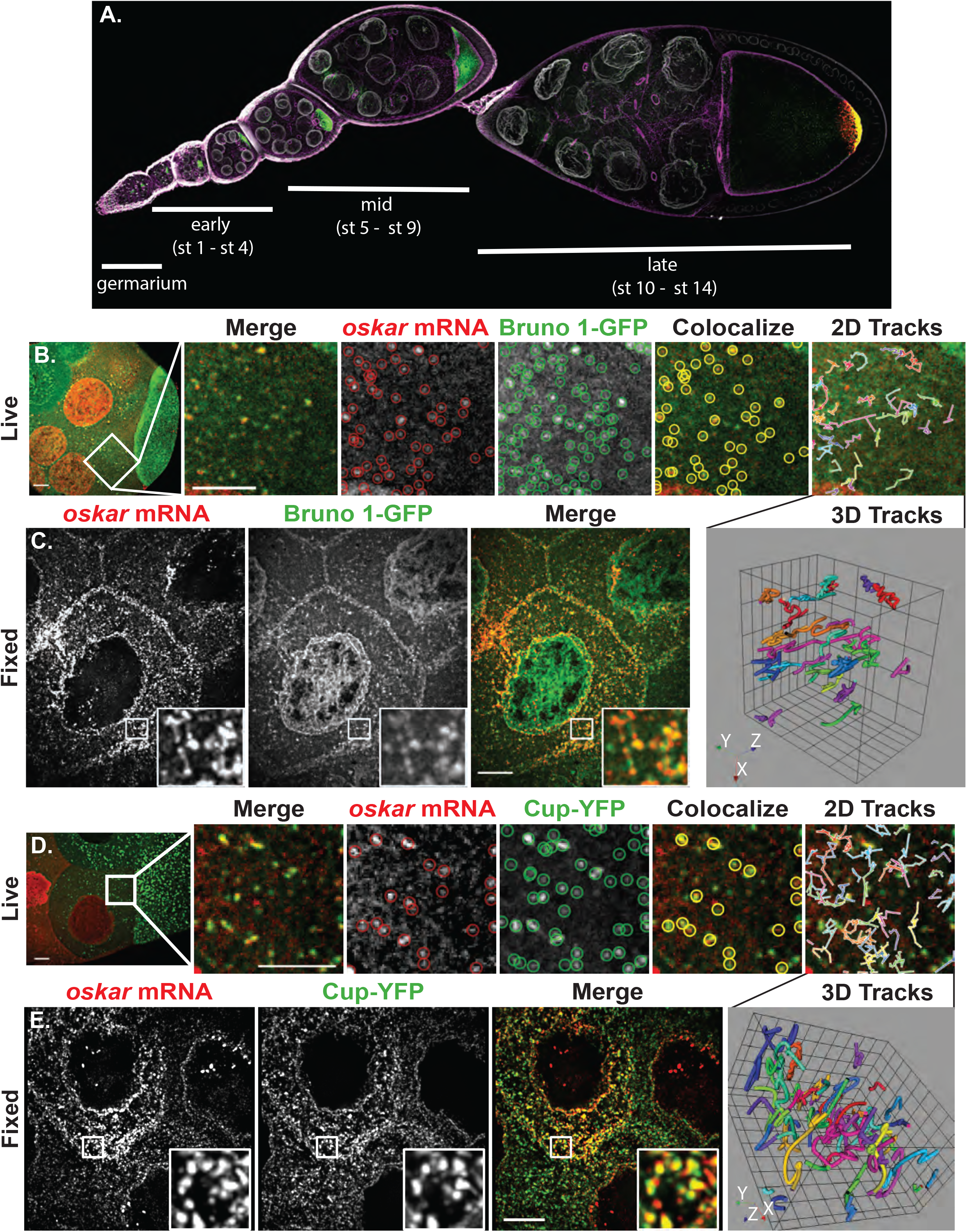
Bru1 and Cup maintain a stable association with *osk* mRNA in the nurse cells and oocyte throughout oogenesis. **(A)** *Drosophila melanogaster* ovariole depicting developmental stages from germarium to stage 10. *osk* mRNA (green), Osk protein (red), F-actin (magenta) and membranes/WGA (white). **(B, C)** Co-visualization of *osk* mRNA and Bru1-GFP in the nurse cells, detected with *osk*-specific molecular beacons (B) or smFISH probes (C). (B) Colocalization analysis in live egg chambers at the 10 min time point of an 18 min XYZCt series taken every 30 sec. Circles indicate *osk* mRNA (red) and Bru1-GFP (green) objects, and colocalization of *osk* mRNA with Bru1-GFP (yellow). Tracked colocalized spots shown as a 2D XYZt-projection or as a 3D representation. (C) Images of a fixed egg chamber acquired at 100x. Images are deconvolved XY max-intensity Z-projections of 7 (live) and 16 (fixed) optical slices (0.3 µm each). **(D, E)** Co-visualization of *osk* mRNA and Cup-YFP in a nurse cell, detected with *osk*-specific molecular beacons (D) or smFISH probes (E). (D) Colocalization analysis in live egg chambers at the 7 min time point of a 20.5 min XYZCt series taken every 30 sec. Circles indicate *osk* mRNA (red) and Cup-YFP (green) objects, and colocalization of *osk* mRNA with Cup-YFP (yellow). Tracked colocalized spots are shown as a 2D XYZt-projection or as a 3D representation. (E) Images of a fixed egg chamber acquired at 100x. Images are deconvolved XY max-intensity Z-projections of 12 (live) and 6 (fixed) optical slices (0.5 and 0.3 µm each), respectively. Scale bars, 10μm.

Bru1’s direct binding to *osk* mRNA has only been demonstrated via biochemical assays (Reveal et al. 2011). We carried out live and fixed-cell experiments to elucidate the spatio-temporal requirements for Bru1 recruitment to *osk* mRNA, using *osk*-specific molecular beacons and smFISH probes, respectively. To define the dynamics of *osk* mRNA and Bru1-GFP association in live egg chambers, we chose to use the previously published, endogenously tagged, homozygous lethal, Bru1-GFP fly line (Nagarkar-Jaiswal et al. 2015, Bose et al. 2022). We also confirmed via immunofluorescence (IF) studies that an anti-Bru1 antibody signal colocalized with the GFP signal (Fig. S1A), indicating that the localization of Bru1-GFP reports on that of endogenous Bru1. We analyzed *osk* mRNA colocalization with Bru1-GFP in both the nurse cells’ cytoplasm and the ooplasm at mid stages after nurse cell microinjections with five different *osk*-specific molecular beacons (Fig. 1B) (Bratu et al. 2003; Bayer et al. 2018). Using the object-based image analysis software Icy (de Chaumont et al. 2012), we found that 57±3% (n=5) of *osk*-containing particles associated with Bru1-GFP in the nurse cells, and 43±4% (n=6) in the oocyte (Figs 1B, S1B). Tracking analyses revealed that colocalized particles traveled together for an average of 8.5±1 min with an average total displacement of 37±5 µm (n = 11), indicating a stable association between *osk* mRNA and Bru1-GFP (2D & 3D tracks in Figs 1B, S1B; Movies S1 and S2). At high magnification, large mRNP particles were observed in fixed egg chambers, indicating the association of *osk* mRNA and Bru1 as multiple copies in both the nurse cell cytoplasm and the oocyte (Figs 1C, S1B). Object-based analysis of large mRNP particles can lead to an underestimation of the degree of colocalization (de Chaumont et al. 2012). Live-cell imaging provided similar colocalization results (45±1%). Together, these two approaches demonstrate a stable association of *osk* mRNP with Bru1 in the nurse cells and the oocyte.

In contrast to Bru1, Cup does not contain known RNA-binding domains, nor is there any evidence for direct binding of Cup to *osk* mRNA. Instead, binding between Cup and *osk* mRNP was proposed to occur through association with other proteins, including Bru1 (Wilhelm et al. 2003; Akira Nakamura, Sato, and Hanyu-Nakamura 2004). However, the precise details of spatial and temporal recruitment of Cup to *osk* mRNP are unresolved. Previous IF experiments indicated that Cup localizes at the germline cell periphery (Keyes and Spradling 1997). However, endogenously tagged Cup-YFP (Lowe et al. 2014) demonstrated the presence of Cup throughout the nurse cells and oocyte cytoplasm in various size condensates, proving once again the limitation of antibody accessibility (Fig. 1D). Colocalization of *osk* mRNA with Cup-YFP in nurse cell cytoplasm averaged 59±2% (n=5) and in the ooplasm 73±3% with as high as 84% (n=5) (Figs 1D, S1C). Colocalization of *osk* mRNA with Cup occurs in a stable complex that travels for over 50±6 µm (average displacement) for an average of 7±1 min (2D & 3D tracks in Figs 1D, S1C; Movies S3, S4). At higher magnification, *osk* mRNA colocalized with Cup-YFP (57±1% n=20) and formed large puncta in the nurse cells’ cytoplasm and ooplasm during mid-oogenesis in fixed egg chambers (Figs 1E, S1C). Together, the image analysis results for live and fixed egg chambers demonstrate that Cup is a core member of the *osk* mRNP complex within the nurse cells’ cytoplasm and ooplasm prior to *osk*’s anchoring at the posterior cortex.

### Bru1 affects Cup protein and *osk* mRNA expression levels, but not their colocalization

It was previously suggested that Bru1 is necessary for recruiting Cup to the *osk* mRNP (Akira Nakamura, Sato, and Hanyu-Nakamura 2004; Wilhelm et al. 2003). To test this hypothesis, we utilized Gal4 drivers under the control of different promoters to identify the most efficient and controlled timing of *bru1* knockdown. In strong Bru1 mutants, it has been shown that Bru1 is crucial for development of the germarium (Webster et al. 1997). Therefore, it was necessary to choose Gal4 drivers that initiate the knockdown of Bru1 after the germarium stage. The *otu-Gal4* driver expresses very low Gal4 levels in the germarium, and development of *bru1*^*RNAi*^ egg chambers arrests around stage 4/5. This driver circumvents the limitations of strong Bru1 mutants, making it ideal for studying Bru1’s role during early oogenesis, after the egg chambers progress through the germarium (Fig. S2A). The efficiency of *bru1* knockdown was confirmed via IF, where a strong Bru1 signal was detected in the germarium (Fig. S2A arrow), with only background-level signal detected in egg chambers starting at stage 1/2 (Fig. S2A).

To assess whether Cup can associate with *osk* mRNP in the absence of Bru1, we generated fly lines expressing Cup-YFP in an *otu-Gal4>bru1*^*RNAi*^ background. For our analysis, we selected egg chambers where Bru1 levels were severely reduced; with such strong depletion of Bru1, a change in the *osk* mRNA/Cup colocalization would be expected if Bru1 were crucial for Cup recruitment to *osk* mRNA. On the contrary, we found that the *osk* mRNA/Cup colocalization percentage was unchanged in egg chambers with *bru1*^*RNAi*^ (37±2%) (Fig. 2B, C) as compared to *wild type* (*wt*) (35±1%) (Fig. 2A, C) backgrounds, indicating that, during early oogenesis, Cup can be recruited to *osk* mRNP via proteins other than Bru1.

**Figure 2.**
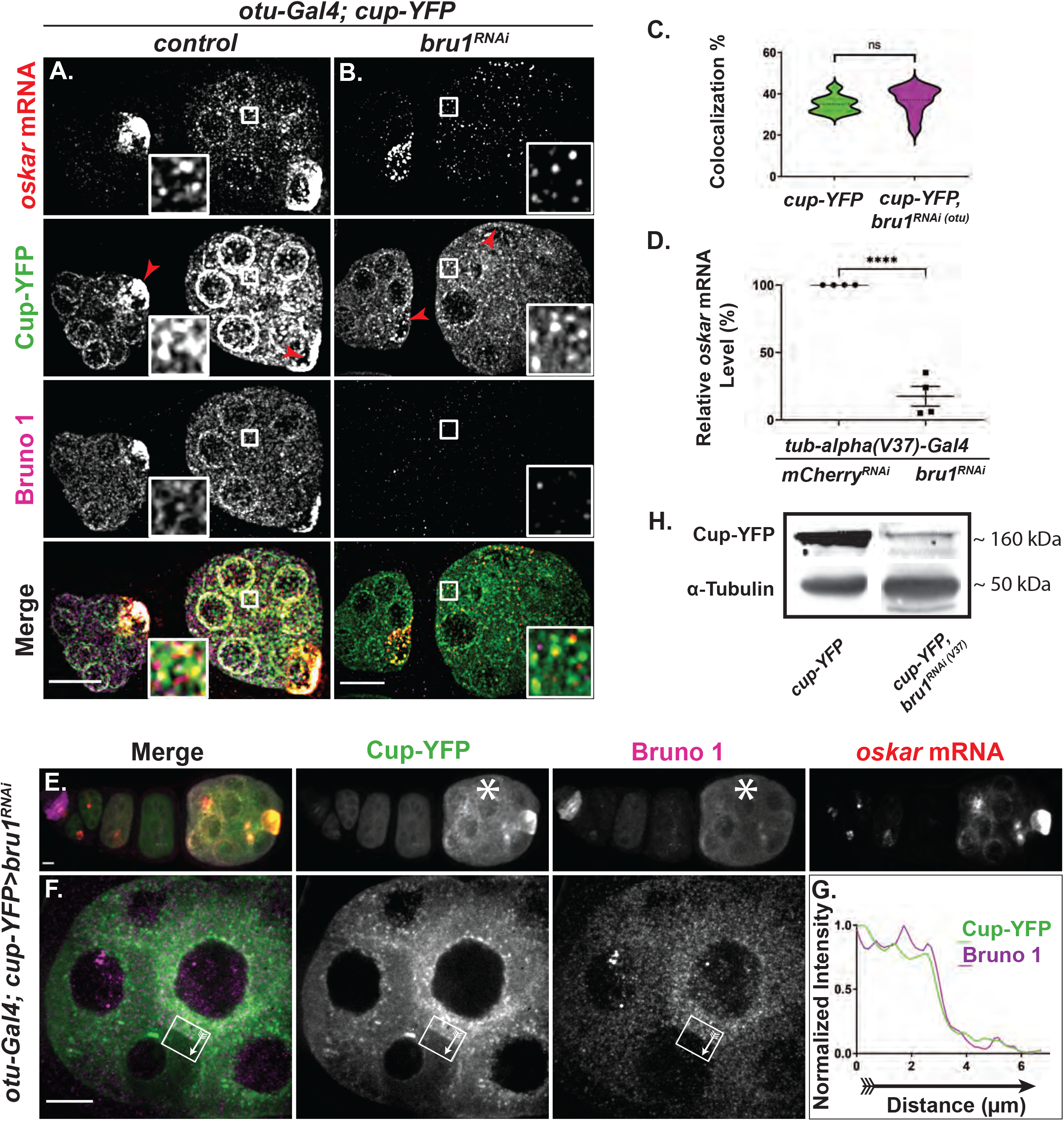
Reduced Bru1 levels affect Cup and *osk* mRNA expression levels, but not their colocalization. **(A, B)** *osk* mRNA and Bru1 co-visualized via smFISH and IF in stage 2 and 3 egg chambers expressing Cup-YFP in *wt* (A) or *bru1*^*RNAi*^ (B) backgrounds. Red arrowhead = oocyte. Images are deconvolved XY max-intensity Z-projections of 17 optical slices (0.3 µm each). **(C)** Colocalization percentage of *osk* mRNA with Cup-YFP in *wt* and *bru1*^*RNAi*^ egg chambers (*wt* n=9, *bru1*^*RNAi*^ n=11, ns = not significant). **(D)** RT-qPCR quantification of endogenous *osk* mRNA in the indicated backgrounds normalized to *rp49* mRNA (mean ± SEM; ****p<0.0001; *mCherry*^*RNAi*^ n=4, *bru1*^*RNAi*^ n=4). **(E-G)** Chain of egg chambers expressing Cup-YFP and various levels of *bru1* knockdown analyzed via IF, with the starred egg chamber showing lower knockdown efficiency and increased Bru1 and Cup-YFP signal (E). Fluorescence intensity analysis of the egg chamber expressing Cup-YFP and different levels of *bru1*^*RNAi*^ in neighboring nurse cells as detected with IF (F). The white arrow inside the box shows the direction of the intensity analysis (G). Images are XY max-intensity Z-projections of 26 (E: 40x) and 18 deconvolved (F: 100x) optical slices (0.3 µm each). **(H)** Immunoblot against GFP and α -Tubulin using lysates prepared from egg chambers expressing Cup-YFP in the indicated backgrounds. Scale bars, 10 μm.

Using *osk* smFISH probes in Bru1-depleted egg chambers, we found that the *osk* mRNA signal was decreased as compared to *wt* (Fig. 2B vs A). We confirmed our visual observations with RT-qPCR analysis. For preparation of whole ovary lysates, we chose a stronger homogeneously expressed Gal4 driver, maternal *tub-alpha* (V37).

This driver allows for germarium development, as it initiates the knockdown at stage 2, and presents full developmental arrest by stage 3/4. We found that in *bru1*^*RNAi*^ egg chambers, *osk* mRNA level was reduced to 18±7% of the level detected in *wt*, indicating that Bru1 is actually necessary for maintaining normal *osk* mRNA levels (Fig. 2D).

Though it was previously reported that *osk* mRNA stability was not mediated by Bru1 through the BRE sites (Kanke et al. 2015), we speculated that in BRE mutants, Bru1 is still recruited to the transcript via Cup, thus maintaining *osk* mRNA stability. Since independent binding of Cup to *osk* mRNA has not yet been demonstrated, it prompted us to search for factors that would recruit Cup to *osk* mRNP. Cup also interacts with eIF4E and Barentsz (Btz), two known components of *osk* mRNP (Wilhelm et al. 2003; Styhler et al. 1998). Systematic knockdowns of paired factors demonstrated that when pairing *bru1*^*RNAi*^ with either *btz*^*RNAi*^ or *eIF4E*^*RNAi*^, egg chambers arrest at a stage similar to *bru1* knockdown alone (Fig. S2D vs A) without any impact on *osk* mRNA/Cup-YFP colocalization observed in *wt* egg chambers [35±1% (*wt* n=9) vs 36±2% (*bru1*^*RNAi*^: *btz*^*RNAi*^ n=6) vs 30±2% (*bru1*^*RNAi*^: *eIF4E*^*RNAi*^ n=5)]. Simultaneous knockdown of *btz* and *eIF4E* arrested egg chamber development mid-oogenesis and had no effect on *osk* mRNA/Cup-YFP colocalization [56±1% (*wt* n=10) vs 59±1% (*btz*^*RNAi*^: *eIF4E*^*RNAi*^ n=7)] (Fig. S2D). These results suggest Cup’s recruitment to *osk* mRNP may be redundant or yet involve other protein factors. An intriguing speculation is that the IDRs of Cup may actually facilitate direct binding to *osk* mRNA, as IDR domains are known to bind mRNAs in other species (Smith et al. 2016). This speculation was supported by results of a recent study that suggest that Cup is able to directly bind to mRNAs without any other factors (Pekovic et al. 2022).

Surprisingly, we also found that Cup-YFP expression was decreased in *bru1*^*RNAi*^ egg chambers and the YFP signal was no longer concentrated in the oocyte (Fig. 2B vs A, arrowheads). This was unexpected, since there were no previous implications for Bru1 in the regulation of Cup expression. To confirm Bru1’s possible role in *cup* gene expression regulation, we exploited the heterogenous levels of *otu-Gal4* expression across egg chambers, where a few egg chambers “escaped” and developed further (Fig. 2E) while in some instances neighboring nurse cells presented varying degrees of the knockdown, thus creating mosaics (Fig. 2F). Convincingly, Cup expression levels directly correlated with those of Bru1. In egg chambers where Gal4 expression led to a weaker *bru1* knockdown and persisting Bru1 signal, Cup-YFP levels were also higher (Fig. 2E, asterisk). Furthermore, in neighboring mosaic nurse cells with different *bru1* knockdown efficiencies, Bru1 and Cup signal intensities remained directly correlated (Figs 2F, G), as did Bru1 and *osk* mRNA signals (Figs S2B, C). Western blot analysis confirmed a severe decrease in Cup protein when Bru1 levels were reduced with the homogeneous expressing Gal4 driver (*Gal4*^*V37*^) (Fig. 2H). Taken together, we unveiled Bru1’s importance for both *osk mRNA* and Cup stability during early-mid oogenesis.

### Cup regulates Bru1 protein expression, distribution, and its association with *osk* mRNP

Bolstered by our findings that Bru1 is not necessary for Cup’s recruitment to *osk* mRNA during early oogenesis, we sought to investigate whether Cup mediates the effects of Bru1 on *osk* mRNA regulation. Cup has been previously shown to interact with numerous proteins and post-transcriptionally regulate multiple mRNAs including *osk* mRNA (Clouse, Ferguson, and Schupbach 2008; Piccioni, Zappavigna, and Verrotti 2005; Piccioni et al. 2009; Verrotti and Wharton 2000; Nakamura, Sato, and Hanyu-Nakamura 2004).

We evaluated the knockdown of *cup* using the *otu-Gal4* driver and found that the expression of Cup is highly reduced in egg chambers beginning at stage 1/2 (Fig. S3A arrow indicates signal in germarium). *cup*^*RNAi*^ egg chambers phenocopy strong *cup* mutants (Keyes and Spradling 1997), developing with almost wild type morphology until stage 5, and rarely progressing past stage 7. In *cup*^*RNAi*^ egg chambers, *osk* mRNA no longer accumulates into large foci, appearing smaller in size in the nurse cell cytoplasm, especially during mid oogenesis (Figs 3A vs B, 3C vs D). We also observed that *osk* mRNA levels were decreased in *cup* knockdowns (reduced to 22±6% of *wt*) (Fig. S3B) just as previous work showed in *cup* mutant egg chambers (Broyer, Monfort, and Wilhelm 2017).

**Figure 3.**
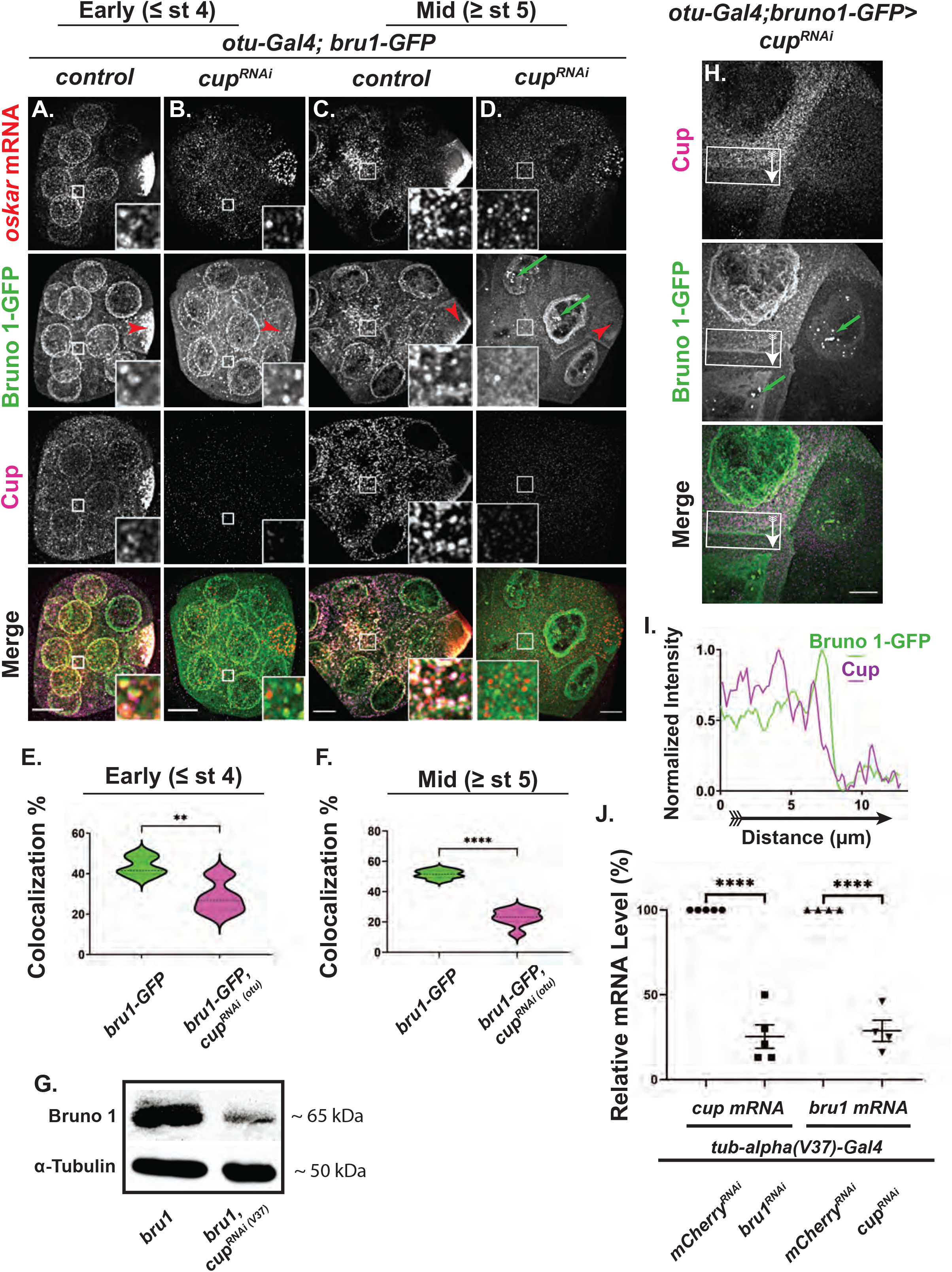
Bru1 protein expression, distribution, and association with *osk* mRNP are regulated by Cup. **(A-D)** *osk* mRNA and Cup detected via smFISH and IF in early and mid-stage egg chambers expressing Bru1-GFP in *wt* or *cup*^*RNAi*^. Early oogenesis, *wt* (A) and *cup*^*RNAi*^ (B). Mid oogenesis, *wt* (C) and *cup*^*RNAi*^ (D). Green arrows highlight the nuclear aggregation of Bru1-GFP in *cup*^*RNAi*^ egg chambers. Red arrowheads = oocyte. Images are deconvolved XY max-intensity Z-projections of 19 (*wt*) and 16 (*cup*^*RNAi*^) optical slices (0.3 µm each) during early, and 22 (*wt*) and 16 (*cup*^*RNAi*^) during mid stage egg chambers. **(E)** Colocalization percentage of *osk* mRNA with Bru1-GFP in *wt* and *cup*^*RNAi*^ back-grounds during early oogenesis (**p<0.0021, ****p<0.0001; *wt* vs *cup*^*RNAi*^: n=5 and n=8). **(F)** Colocalization percentage of *osk* mRNA with Bru1-GFP in *wt* and *cup*^*RNAi*^ back-grounds during mid oogenesis (**p<0.0021, ****p<0.0001; *wt* vs *cup*^*RNAi*^: n=6 and n=8). **(G)** Immunoblot against Bru1 and α -Tubulin using lysates prepared from egg chambers in indicated backgrounds. **(H, I)** Fluorescence intensity analysis of an egg chamber expressing Bru1-GFP and different levels of *cup*^*RNAi*^ knockdown in neighboring nurse cells, detected with IF. Green arrows highlight nuclear aggregation of Bru1-GFP (G). The white arrow inside the box shows the direction of the intensity analysis (H). Images are deconvolved XY max-intensity Z-projections 16 optical slices (0.3 µm each). **(J)** RT-qPCR quantification of endogenous *cup* mRNA and *bru1* mRNA in the indicated backgrounds normalized to *rp49* mRNA (mean ± SEM; ****p<0.0001; *mCherry*^*RNAi*^ n=5, *bru1*^*RNAi*^ n=5 *mCherry*^*RNAi*^ n=4, *cup*^*RNAi*^ n=4). Scale bars, 10 μm.

Until stage 3 of oogenesis, Bru1-GFP signal became slightly more diffused in f54the nurse cells, but its intensity remained comparable to *wt* (Figs 3B vs A), however it no longer concentrated in the oocyte (Figs 3B vs A, D vs C, red arrowheads). Beginning around stage 4 and persisting during mid-oogenesis, Bru1-GFP expression became highly diffused and no longer formed large puncta in the nurse cells’ cytoplasm (Figs 3D vs C). Intriguingly, Bru1-GFP formed extremely large and highly concentrated puncta in the nurse cell nuclei that did not include *osk* mRNA (Figs 3D, H, S3C, green arrows). We confirmed the nuclear and cytoplasmic localization of endogenous Bru1 in *cup*^*RNAi*^ background via IF in order to ensure that the endogenous protein displays the same phenotype as the GFP-tagged protein (Figs 4C, E, S3D, S4A). When the full volume of the nurse cell nuclei was visualized, the large nuclear aggregation phenotype was highly penetrant with either Bru1 antibody (Figs 4C, E, green arrows) or Bru1-GFP (Figs 3D, G, S3C, green arrows).

**Figure 4.**
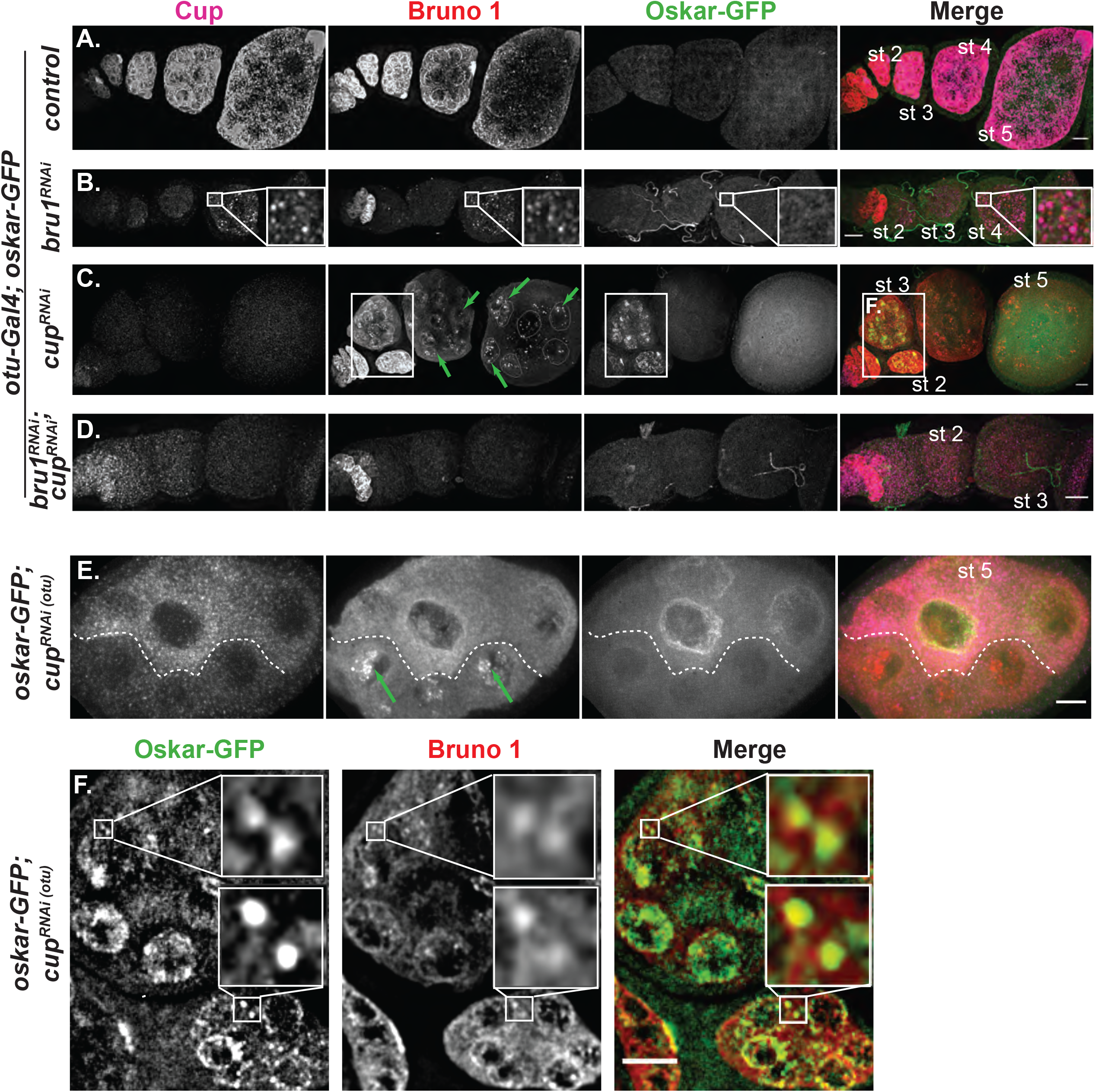
Cup directs translational silencing, while Bru1 stimulates translational activation of *osk* mRNA during early stages of oogenesis. **(A-D)** Cup and Bru1 visualized via IF in egg chambers expressing Osk-GFP in the indicated RNAi backgrounds, *wt* (A), *bru1*^*RNAi*^ (B), *cup*^*RNAi*^ (C), and *bru1*^*RNAi*^*;cup*^*RNAi*^ (D). Green arrows highlight nuclear aggregation of Bru1; some of the nuclei were outlined for clarity. Images are deconvolved XY max-intensity Z-projections of 29 (*wt*), 30 (*cup*^*RNAi*^), 27 (*bru1*^*RNAi*^) and 34 (*bru1*^*RNAi*^*;cup*^*RNAi*^) optical slices (0.3 µm each). **(E)** Nurse cells expressing Osk-GFP with different levels of *cup*^*RNAi*^ knockdown in neighboring nurse cells delineated by a dashed line, detected with IF. Green arrows highlight nuclear aggregation of Bru1. Image is XY max-intensity Z-projection of 34 optical slices (0.3 µm each). **(F)** Inset from (C) showing premature Osk-GFP and Bru1 colocalization. Scale bars, 10 μm.

Since Bru1’s level and cytoplasmic localization were altered in *cup*^*RNAi*^ egg chambers, we analyzed the colocalization of *osk* mRNA with Bru1-GFP. In early stages, *osk* mRNA colocalization with Bru1-GFP only moderately decreased (44±2% vs 28±3%) (Fig. 3E); however, in stage 5 or older egg chambers, the colocalization was strongly reduced (52±1% vs 23±2%) (Fig. 3F). When comparing these results with our live cell imaging data, we observed similar distribution patterns (Movie S5). Taken together, these data indicate that Cup plays a crucial role in the proper formation of *osk* mRNP, and in the proper localization of Bru1.

The diffused Bru1 expression phenotype raises the question of whether the decrease in Bru1-GFP signal intensity represents a decrease in total Bru1 protein or merely its redistribution in the egg chamber, as previously suggested (Broyer, Monfort, and Wilhelm 2017). To test this possibility, we carried out Western blot analysis for endogenous Bru1 in *cup*^*RNAi*^, using the stronger/homogeneous Gal4 driver (*Gal4*^*V37*^), which led to an early developmental arrest (∼stage 5/6) (Fig. 3G). Our results do not echo those from previous Western blot analysis of *cup* mutant egg chambers, where no decrease in Bru1 levels was detected. However, similar to our results from IF experiments, they saw a comparable reduction of Bru1 signal in the oocyte during early stages, contradicting their Western blot results (Broyer, Monfort, and Wilhelm 2017).

Due to the experimental differences (i.e. the use of *cup* mutant egg chambers and different antibodies for the Western blot analysis), it is difficult to compare and contrast the results of that study with those of ours. We confirmed our Western blot analysis with anti-GFP antibody, where we detected the same decrease in Bru1 levels when *cup* expression was knocked down with the *otu-Gal4* driver. We also detected two Bru1-GFP bands, indicating a possible post-translational modification of Bru1, in agreement with a previously proposed hypothesis (Fig. S3E) (M. Snee et al. 2008). To further confirm that Bru1-GFP levels indeed decreased in *cup*^*RNAi*^ egg chambers, we analyzed Bru1-GFP levels in mosaic *cup*^*RNAi*^ egg chambers and found that nurse cells with a lower intensity Cup signal also showed a decreased Bru1-GFP signal in the cytoplasm, with large, concentrated puncta visible in the nuclei (Figs 3H, I, green arrows). We found a similar reduction of endogenous Bru1 between mosaic nurse cells in *cup*^*RNAi*^ background (Fig S3D red arrow). Our results illuminate a new and critical role for Cup in maintaining normal Bru1 expression during early oogenesis.

We decided to further investigate whether Cup regulates Bru1 at the protein or mRNA level. Interestingly, we found that when Cup levels were reduced, *bru1* mRNA levels were significantly decreased (Fig. 3J), bringing credence to our Western blot results. Moreover, we also demonstrated that Bru1 is necessary to maintain *cup’s* normal mRNA levels (Fig. 3J). Remarkably, this indicates a surprisingly reciprocal regulation where Cup and Bru1 control each other’s expression at the mRNA level.

We used the RBPmap web server to identify numerous candidates that could mediate this mutual regulation (Paz et al. 2014). Analysis of *bru1* mRNA shows that multiple proteins have putative binding sites within the sequence, including Bru1 itself, possibly recruiting Cup to *bru1* mRNA (Paz et al. 2014). Most interestingly, we also found that the *cup* mRNA sequence contains numerous BRE sites, which may enable direct binding of Bru1 to the *cup* transcript. This association could result in the stabilization of *cup* mRNA, perhaps through Bru1 recruiting Cup itself. While this is a plausible explanation, the Rangan group recently showed that Bru1 is necessary for translational repression of *pgc* mRNA (Flora et al. 2018). PGC directs global transcriptional silencing by inhibiting RNA polymerase II activity. Conceivably, the decreased levels of *cup* and *osk* mRNAs may be due to this indirect effect of Bru1. These two possible mechanisms of *cup* mRNA regulation via Bru1 are not mutually exclusive and require further exploration.

### Cup directs translational silencing during early oogenesis, while Bru1 stimulates translational activation of *osk* mRNA

Cup’s role in the translational repression of *osk* mRNA has only been established in the oocyte during mid to late oogenesis (Styhler et al. 1998). Premature Osk protein expression in *bru1* mutants has never been detected, with Bru1’s role in translational repression only demonstrated using transgenes and ovary lysates (Castagnetti et al. 2000; Chekulaeva, Hentze, and Ephrussi 2006; Kim-Ha, Kerr, and Macdonald 1995; Reveal et al. 2010; Reveal et al. 2011). We aimed to clarify the roles of both factors in the translational regulation of *osk* mRNA during early stages of oogenesis by using egg chambers that express endogenously tagged Osk-GFP. We confirmed that Osk-GFP mimics the wild type protein expression via an anti-Osk antibody (Figs S4D, E). Surprisingly, both Osk-GFP and the anti-Osk antibody enabled us to detect very low Osk levels during earlier stages, which were visualized as colocalized signals (Figs S4G, H). Since such a premature and ectopic Osk signal has never been reported, it is possible that it was interpreted to be nonspecific due to the quality of the Osk antibodies, as well as the reduced resolution of the detection approaches. This presents a new idea of a possible “leaky” control of *osk* mRNA translational repression. While the antibody is reliable for detecting Osk protein, it generates a strong background signal, especially during earlier stages of oogenesis where we focused our study, therefore we proceeded using the tagged protein for our analysis. Once again, we chose the *otu-Gal4* driver to minimize the knockdown of Bru1 in the germarium, allowing us to only address the early stages of oogenesis.

To our surprise, premature Osk-GFP was not detected when Bru1 levels were decreased in early-stage egg chambers, an indication that *osk* mRNA translation remained repressed. Cup expression was reduced in these egg chambers, but it nevertheless persisted and remained colocalized with *osk* mRNA even when Bru1 levels were no longer detectable (Figs 2B vs A, 2C). In egg chambers with less efficient *bru1* knockdown, Cup and Bru1 were still colocalized (Fig. 4B inset). Altogether, this suggests that Cup is sufficient to maintain the translational repression of *osk* mRNA (Fig. 4B). Confirming this finding, we could not detect premature Osk-GFP in a different *bru1*^*RNAi*^ line, as well as in two *bru1* mutant and deficiency fly lines (data not shown). Nonetheless, this does not exclude the possibility that Bru1 may be present below our detection limits and could facilitate sufficient translational repression.

By contrast, *cup* knockdown led to the accumulation of a strong Osk-GFP signal in the nurse cells, particularly within the nuclei and their periphery, as well as within the oocyte (Figs 4C, S4A, S4B). We validated the observed premature Osk-GFP expression by crossing a combination of four different *cup* mutant fly lines (i.e., *cup*^*1/15*^, Fig. S4C). In *cup*^*RNAi*^ egg chambers, the Osk-GFP signal was detected as early as stage 1. The strongest Osk-GFP signal was surprisingly observed in stages 2-3, when Bru1 expression was still comparable to *wt*, confirming that Bru1 alone, without Cup, is not sufficient to maintain the translational repression of *osk* mRNA during these early stages (Fig. 4C, 4F, S4A). Beginning at stage 5, Osk-GFP signal was drastically reduced in *cup*^*RNAi*^ egg chambers, becoming undetectable by stage 7. Interestingly, these are the stages when Bru1-GFP aggregated in the nurse cell nuclei and no longer colocalized with *osk* mRNA in the cytoplasm (Figs 4C, E, arrows), implying that Bru1 plays a key role in activating *osk* mRNA translation.

To further test this idea, we postulated that if Bru1 plays a crucial role in activating *osk* mRNA translation, a double knockdown of *cup* and *bru1* will significantly reduce any activation that Bru1 mediates in *cup*^*RNAi*^ egg chambers. However, if Bru1 serves only to help repress translation, without a role in activating translation, the double knockdown would present premature Osk-GFP expression due to the simultaneously reduced Cup levels. Remarkably, we did not observe any premature Osk-GFP in these egg chambers (Fig. 4D), even though *osk* mRNA was still detected, albeit at a significantly reduced level (Fig. S4A). To further confirm Bru1’s role in activating *osk* mRNA premature translation, we again exploited the differences in *cup*^*RNAi*^ knockdown efficiency in neighboring nurse cells. Bru1 formed large puncta in the nurse cell nuclei (Fig. 4E, arrows) and Osk-GFP was undetectable when the levels of Cup and Bru1 signals were strongly reduced as compared to *wt*. However, we observed a strong premature Osk-GFP expression in nurse cells where both Cup and Bru1 were detectable but at very low levels (Fig. 4E). This indicates that Cup must persist above a certain threshold level to maintain translational repression, and similarly there is a threshold level for Bru1 to facilitate translation. Additionally, Bru1 and Osk-GFP colocalized in the nurse cell cytoplasm in large puncta indicating possible translational hubs of *osk* mRNA via Bru1 (Fig. 4F). We confirmed these results by detecting endogenous Osk in the presence or absence of the Osk-GFP (Figs S4, FG). While the Osk signal reported by the antibody was difficult to detect, we used an advanced image acquisition set up, optimized to resolve low signals.

### Cup facilitates the association of osk mRNA with P-bodies and is necessary for proper P-body morphology

Knowing that *osk* mRNA localizes into P-bodies, where translationally repressed mRNAs are stored (Fan, Marchand, and Ephrussi 2011; A. Nakamura et al. 2001), we hypothesized that with reduced levels of Cup, *osk* mRNA would not associate with P-bodies, thus leading to *osk*’s premature and ectopic translation and degradation. To test this notion directly, we examined the recruitment of *osk* mRNA into P-bodies, using the P-body marker Me31B-GFP. First, we determined that Bru1, Cup, Me31B-GFP and *osk* mRNA all associate in the cytoplasm, forming large aggregates in *wt* egg chambers, indicating that they are components of the same condensate (Fig. S5A). As *osk* mRNA accumulates in a tight crescent at the oocyte’s posterior pole, it becomes poised for translation. During this time, *osk* mRNA signal highly exceeds the expression signal of Me31B and Cup, indicating a reorganization of the components of these condensates, with a likely dissociation of *osk* mRNA from P-bodies (Fig. S5B).

Next, we generated flies co-expressing the *otu-Gal4* driver and endogenously tagged Me31B-GFP in combination with *bru1*^*RNAi*^ or *cup*^*RNAi*^. In *bru1*^*RNAi*^ egg chambers, we did not detect a change in the association of *osk* mRNA with Me31B-GFP, indicating that *osk* mRNA continues to localize into P-bodies, and it is translationally repressed when Bru1 levels were reduced (Fig. S5C, D). However, in *cup*^*RNAi*^ background, though Me31B-GFP was reduced, a considerable amount of signal persisted throughout the egg chambers (Fig. 5 A vs B and F vs G). This was not surprising, as Cup has been shown to be partially important for maintaining Me31B levels (Broyer, Monfort, and Wilhelm 2017). Me31B-GFP formed condensates in early stages of *cup*^*RNAi*^ egg chambers, but the colocalization of *osk* mRNA with Me31B-GFP was decreased (49±1% in *wt* to 43±1% in *cup*^*RNAi*^) (Figs 5A vs B, C), however the dissociation of *osk* mRNA from Me31B-GFP became more significant during mid-oogenesis (51±1% in *wt* to 40±1% in *cup*^*RNAi*^) (Figs 5F vs G, H). The decreased percentage in their colocalization becomes even more significant when taking into account the already reduced levels of *osk* mRNA in *cup*^*RNAi*^ egg chambers. If the association of *osk* mRNA with P-bodies were not mediated via Cup, these lower levels of *osk* mRNA would instead increase the percentage of colocalization with Me31B, as transcripts within P-bodies are protected from degradation. Our data indicates that Cup is necessary for *osk* mRNA’s association with P-bodies, implicating that when *osk* mRNA is not housed in P-bodies, translational repression and mRNA stability are compromised.

**Figure 5.**
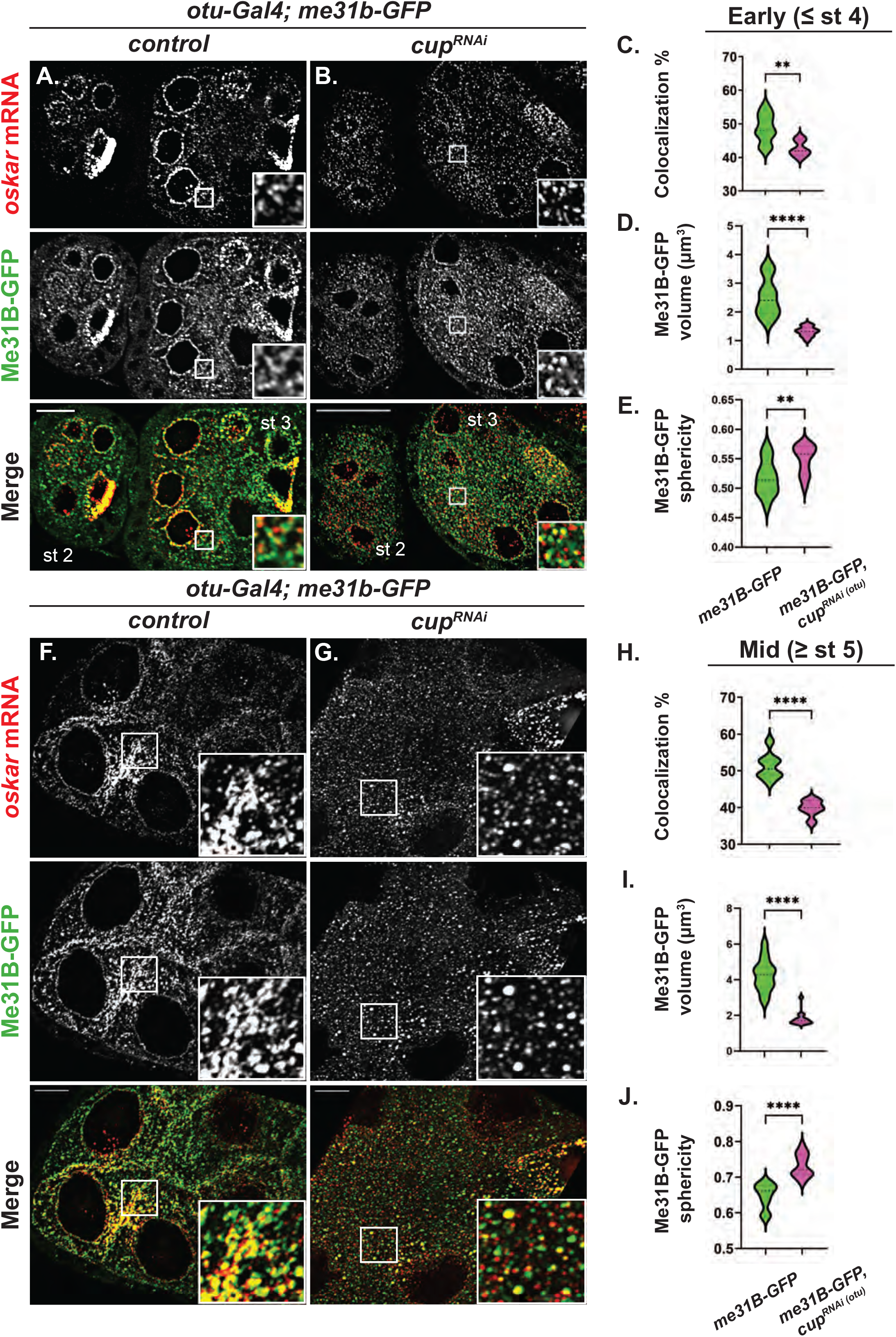
Cup facilitates the recruitment of *osk* mRNA into P-bodies and regulates the condensates’ physical state. **(A, B, F, G)** Representative images of *osk* smFISH experiments expressing Me31B-GFP in *wt* and *cup*^*RNAi*^ backgrounds in early stage (A, B) and mid stage (F, G) egg chambers. Images are deconvolved XY max-intensity Z-projections of early stage: 26 (*wt*), 25 (*cup*^*RNAi*^), 25 (*bru1*^*RNAi*^) and mid stage: 31 (*wt*), 25 (*cup*^*RNAi*^) egg chambers, optical slices (0.3 µm each). **(C, H)** Colocalization quantification of *osk* mRNA with Me31B-GFP was performed in *wt* and *cup*^*RNAi*^ backgrounds in early stage (**p=0.0076; *wt* vs *cup*^*RNAi*^: n=9 and n=6) (C) and mid stage (****p<0.0001; *wt* vs *cup*^*RNAi*^: n=8 and n=6) (H) egg chambers. **(D, I)** Volume analysis of Me31B-GFP condensates was performed in *wt* and *cup*^*RNAi*^ backgrounds in early stage (****p<0.0001; *wt* vs *cup*^*RNAi*^: n=13 and n=13) (D) and mid stage (****p<0.0001; *wt* vs *cup*^*RNAi*^: n=14 and n=14) (I) egg chambers. **(E, J)** Sphericity analysis of Me31B-GFP condensates was performed in *wt* and *cup*^*RNAi*^ backgrounds in early stage (**p=0.0052; *wt* vs *cup*^*RNAi*^: n=14 and n=14) (E) and mid stage (****p<0.0001; *wt* vs *cup*^*RNAi*^: n=14 and n=14) (J) egg chambers. Scale bars, 10 μm.

Previous work from multiple groups have shown that the physical properties of Me31B condensates correspond to the fate of mRNAs that are regulated in P-bodies in mature oocytes and in early embryos (Sankaranarayanan et al. 2021; Bose et al. 2022). When P-bodies are in a dynamic state, characterized by a smaller and more spherical morphology, mRNAs are released from and/or fail to incorporate into P-bodies. Conversely, the more amorphous, arrested physical state, preceded by a short-lived liquid state, facilitates mRNA storage and the incorporation of mRNAs and proteins into P-bodies (Bose et al. 2022; Sankaranarayanan et al. 2021). We wished to assess the role of Cup in modulating the physical state of P-bodies during oogenesis by analyzing the size and sphericity of Me31B-GFP condensates. In *cup* knockdown, we found a significant decrease in P-body size (early: 2.51±0.18 µm^3^ in *wt* to 1.32±0.05 µm^3^ in *cup*^*RNAi*^; mid: 4.27±0.24 µm^3^ in *wt* to 1.81±0.10 µm^3^ in *cup*^*RNAi*^) (Figs 5D, I). More importantly, the P-body surface also changed significantly from an amorphous to a more spherical morphology (early: 0.52±0.008 µm^3^ in *wt* to 0.55±0.007 µm^3^ in *cup*^*RNAi*^; mid:0.65±0.009 µm^3^ in *wt* to 0.73±0.009 µm^3^ in *cup*^*RNAi*^) (Figs 5E, J). Taken together, these results indicate that Cup is important for modulating the physical state of P-bodies, which in turn facilitates proper storage of translationally repressed mRNAs.

## DISCUSSION

Bru1 and Cup are critical for *osk*’s mRNA life cycle during oogenesis. Here we demonstrate that during early oogenesis when Cup expression level is reduced, Bru1’s association with *osk* mRNA is compromised. This is an important and unexpected result since Bru1’s binding at the BRE sites of *osk* 3’UTR was observed to occur without a binding partner (Reveal et al. 2011). In support of our finding, a recent study demonstrated that the deletion of the N-terminal (Cup-binding) domain of Bru1, that left the RRMs domains intact, led to a largely diffuse Bru1 expression and reduced *osk* mRNP granule assembly (Bose et al. 2022). We believe that both the direct Bru1 binding, and a Cup-stabilized interaction with *osk* mRNA contribute to the formation of a mature *osk* mRNP.

In *cup*^*RNAi*^ throughout early oogenesis (stage 2-4), we observed a robust premature Osk expression despite Bru1 levels persisting, which precludes the notion that small amounts of Bru1 alone can maintain translational repression. This indicates that Bru1 was insufficient for silencing *osk* mRNA translation, even though Bru1 and *osk* mRNA remained colocalized, albeit at a reduced level. In stark contrast, *bru1* knockdown egg chambers did not express Osk prematurely. Cup levels decreased, but the remaining Cup continued to colocalize with *osk* mRNA, sufficiently maintaining its translationally repressed state. It was previously found that premature Osk expression occurred exclusively in the oocyte in egg chambers with deleted eIF4E-binding sites in Cup (Nakamura, Sato, and Hanyu-Nakamura 2004). Therefore all together, it suggests that the Bru1-Cup-eIF4E complex is mostly important for *osk*’s translational repression after the mRNP is deposited into the oocyte, while in the nurse cells, Cup is the essential factor for *osk*’s translational repression.

We further speculate that Bru1 is not only necessary for translation activation of *osk* mRNA at the posterior cortex of the oocyte, but the loss of Cup from the *osk* mRNP allows Bru1 to play its role in translation throughout the egg chamber. For such translation to occur, Bru1 expression must attain a sufficient threshold level which was indicated by using the double knock down of *cup*^*RNAi*^*bru1*^*RNAi*^ and through our mosaic egg chamber analysis. We postulate that the translation activation is facilitated via the N-terminal domain, since Osk protein was not detected in N-terminal-deleted *bru1* mutant egg chambers (Bose et al. 2022). As Cup normally binds Bru1 through this domain, future studies could explore whether other protein factors are recruited to this domain when Cup is removed, thus activating *osk* mRNA translation.

We also observed a previously undescribed phenotype in *cup*^*RNAi*^ egg chambers: Bru1 forms large punctas in nurse cell nuclei devoid of *osk* mRNA. We propose two possibilities: (1) since Cup has been shown to interact with members of the nuclear pore complex (Grimaldi et al. 2007), the nuclear export process is affected leading to an increased concentration of Bru1 in the nuclei which, in turn, would lead to a concentration-dependent aggregation, or (2) as previous *in vitro* studies showed that the site for Bru1 dimerization is the same as that for binding to Cup, the presence of Cup directly blocks Bru1 dimerization (Kim et al. 2015). We can thus postulate that without Cup, Bru1 forms large aggregates due to increased Bru1-Bru1 association in the nucleus, and a subsequent alteration of its normal association with *osk* mRNP. In the nurse cell cytoplasm, for both possibilities, Bru1 is more diffused leading to a lower local concentration thus making aggregation less likely. We favor the second possibility since the N-terminal-deleted Bru1 protein mutants were also diffused in the cytoplasm (Bose et al. 2022).

Collectively, we propose that Bru1 and Cup are in a reciprocal relationship to maintain their proper protein levels and distribution in the egg chamber. This lethality-associated sensitive protein threshold relationship, controlled via auto and/or cross regulation, is widespread in cell biology and has been demonstrated specifically for other RNA binding proteins, such as ELAV in neurons, and for RBPs that regulate translation and RNA integrity (Borgeson and Samson 2005; Pullmann et al. 2007; Samson 1998). Deciphering how expression and function of these CELF family member proteins are regulated is crucial for our understanding of health and disease progression (reviewed in (Ladd 2013; Nasiri-Aghdam, Garcia-Garduño, and Jave-Suárez 2021)).

Intriguingly, Cup’s role in Bru1’s expression extends beyond its protein localization as we found that Cup regulates *bru1* mRNA levels and surprisingly, Bru1 regulates *cup* mRNA levels, possibly creating a feedback loop. Recent work showed that the recruitment and release of mRNAs into/from P-bodies is determined by the physical state of the P-bodies, as well as their integrity (Sankaranarayanan et al. 2021). It was demonstrated that P-bodies are dynamic in size and shape and can progress to smaller, spherical morphologies. Cup has been known to interact with multiple key P-body factors such as Me31B and Tral. In the absence of Cup, during early oogenesis, that P-body morphology is altered and association of *osk* mRNA with Me31B is decreased. At these early stages, we observed the highest premature expression of Osk protein. We also speculate, and a future study could determine, that *bru1*, just like *osk* transcript, is stabilized in P-bodies.

As oogenesis progresses, in *cup* knockdowns, the physical state of P-bodies continues to change, further reducing in volume and becoming more spherical. In this state, the majority of the *osk* mRNA is not associated with P-bodies. The smaller and more spherical morphology of Me31B condensates is also reminiscent of the physical state of P-bodies in the early embryo, where the mRNAs are released for translation. Another study has shown that during early embryogenesis, Cup levels decrease drastically leading to a change in Me31B’s role in the regulation of mRNAs. During this period, mRNA association with Me31B switches from mRNA storage to mRNA degradation (Wang et al. 2017). We also find that *osk* mRNA is unstable when its association with P-bodies is compromised. Our results suggest that Cup dictates the formation of a more amorphous, arrested state of P-bodies. In Cup’s absence, Me31B condensates become more spherical, resulting in *osk* mRNA degradation. However, we cannot rule out the possibility that when Cup is depleted, Me31B levels decrease to such a degree that it can no longer form proper condensates. Nevertheless, we show that Cup is crucial for ensuring proper P-body morphology. Whether the premature *osk* mRNA translation is due to the mRNA’s inability to be recruited into P-bodies or due to its premature release from P-bodies, are two non-mutually exclusive possibilities that warrant further investigation. Moreover, we argue that *osk* mRNA storage is facilitated by the amorphous P-body state, shedding new light on mechanisms of maternal mRNA storage.

Here, we put forth a functional model where Bru1 and Cup, like “yin and yang” partners, play a reciprocal role in supporting each other’s proper expression by maintaining their sensitive protein threshold levels and promoting each other’s wild type localization (Fig 6). The Bru1-Cup complex is necessary for preserving a stable and translationally silent *osk* mRNA, and it plays a crucial role in the association of the *osk* mRNP with P-bodies, thus leading to *osk*’s translational repression. Throughout oogenesis, the physical properties of P-bodies are regulated and modulated by the presence or absence of Cup. In its presence, translationally repressed *osk* mRNA is stored in P-bodies in an arrested state. Upon developmental cues, the mRNA must be released from P-bodies, a process which is facilitated by decreased levels of Cup in the mature egg chamber and early embryo. The resulting change in P-body morphology causes the dissociation of mRNAs from P-bodies which in turn leads to mRNA translational derepression and mRNA degradation. As Cup is removed from the mRNP complex, it may allow for another Bru1-binding factor to take its place, and stimulate translation of *osk* mRNA.

**Figure 6.**
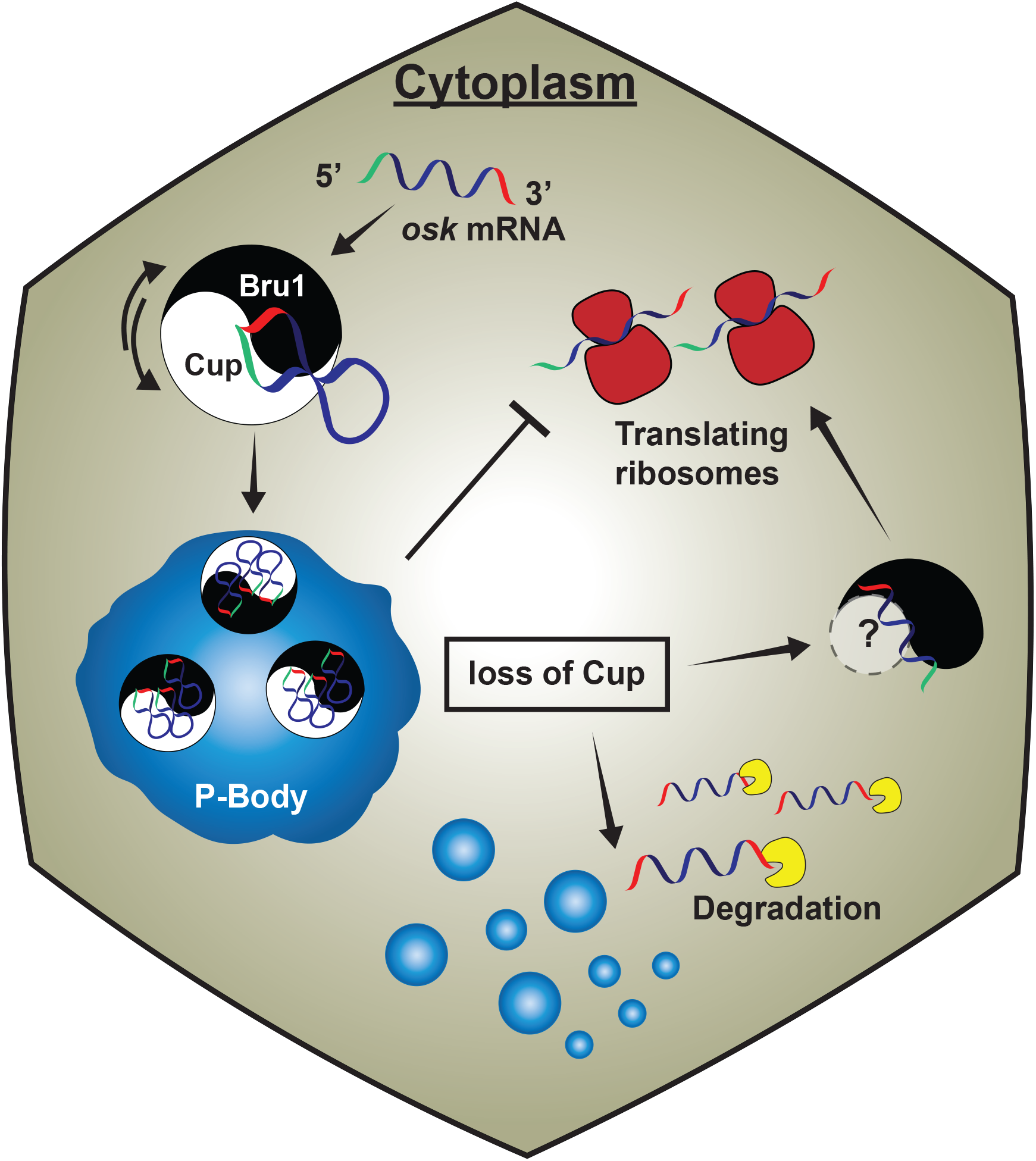
Cup models P-body physical states and governs *osk* mRNA regulation during *Drosophila melanogaster* oogenesis. An intertwined mechanism of post-transcriptional gene regulation during oogenesis, in which a feedback mechanism between Bru1 and Cup maintains a sensitive threshold of each other’s gene expression and distribution. Bru1 and Cup form a complex that associates with *osk* mRNA in the cytoplasm. Cup then links *osk* mRNA to P-bodies, where it is maintained as a stable and translationally silent transcript. Upon loss of Cup, the physical state of P-bodies changes, *osk* mRNP no longer associates with P-bodies that leads to *osk* mRNA translation and ultimate degradation. Translation is facilitated by Bru1 that may occur by recruiting another Bru1-binding factor.

This type of mRNA regulation is conserved beyond *Drosophila* oogenesis, such as in neuronal development, where both mRNA binding proteins and mRNAs stored in phase-separated condensates are necessary for proper memory formation (Tsang et al. 2019) (reviewed in (Arkov 2022)). Here we show, for the first time, that a P-body component regulates the expression of an RNA-binding protein, suggesting that such a mechanism is broadly present in systems where spatio-temporal cues dictate mRNA translation.

## ACKNOWLEDGEMENTS

We thank Dr. A. Spradling (Carnegie Institution for Science), Dr. A. Nakamura (Riken Center for Developmental Biology), Dr. P. Lasko (McGill University) and Dr. M. Lilly (NIH) for their kind gifts of antibodies. We extend a special thanks to Dr. G.B. Gonsalvez (Augusta University) for allowing us to use, and be first to publish, the CRISPR *osk-GFP* fly strain made in his laboratory. We thank the BDSC Indiana and Kyoto DGRC for providing the fly stocks for our study, and the TRiP at Harvard Medical School (NIH/NIGMS R01-GM084947) for providing the transgenic RNAi fly stocks. We thank E. Bagaeva for assistance with fly husbandry and maintenance of stocks, and Dr. S.A.E. Marras (Public Health Research Institute Center, Rutgers University) for the synthesis, labeling and purification of *osk* molecular beacons, and labeling of the *osk* smFISH probes. We thank the Bioimaging Facility at Hunter College for access to the Leica TCS SP8, Nikon spinning disc microscope, AutoquantTCS and Imaris 9.7 software used for image analysis. We thank Dr. P. Feinstein for allowing us to use the Roche Lightcycler instrument. We are grateful to Dr. J.M. McLaughlin and O.S. Omar for their critical comments and especially to Dr. R. Persell for his editorial advice on the manuscript. This work was supported by National Science Foundation (1149738, 1919829), National Institute of Health (1SC1GM135132-01) and Professional Staff Congress-CUNY awards to D.P.B., and a Dissertation Fellowship to L.V.B.

## MATERIALS AND METHODS

### Fly husbandry

All fruit fly stocks were maintained on standard cornmeal agar food at 25°C. Prior to dissection, female flies were fed yeast paste in grape vials for 2-5 days. Fly stocks were obtained from Bloomington *Drosophila* Stock Center: Gal4 inducible TRiP lines: *mCherry* (BL #35785), *cup* (BL #35406), *bru1* (BL #35394 and BL #54812), *btz* (BL #58353), *eIF4E* (BL #34096). Mutant Lines: *cup*^*15*^ (BL #29718) *cup*^*1*^ (BL #4978). Maternal Gal4 driver lines: BL #7063 (*tub-alpha(V37)-Gal4)* and BL #58424 (*otu*). Fluorescently tagged: *Bru1-GFP* (BL #60144) and *Me31B-GFP* (BL #51530). We acquired *Cup-YFP* (DGRC 115-161) from the Kyoto *Drosophila ‘*Stock Center. CRISPR *Oskar-GFP* fly line was a generous gift from Dr. G. Gonsalvez (Augusta University). Experiments were carried out with the *otu-Gal4* driver, unless otherwise indicated.

### Single-molecule RNA FISH (smFISH) and combined smFISH-Immunofluorescence (IF)

smFISH and smFISH-IF were performed as previously described (Bayer et al, 2015). Briefly, ovaries were dissected in Robb’s medium (Recipe, 2011), fixed in 4% PFA in 1X oocyte buffer for 10 min. After fixation, egg chambers were pre-hybridized in wash buffer (2X SSC, 10% formamide), incubated with the *osk*-CY5 probes (Supplemental_Methods_S1) overnight at 37°C, followed by washing with wash buffer and mounting with ProLong™ Diamond Antifade Mountant (Life Technologies) on a glass slide using a #1.5 cover glass. Combined smFISH-IF was carried out with the following modification: after fixation, the egg chambers were permeabilized for 2 h in 1% Triton X-100 in 2X SSC with 1% BSA followed by incubation with *osk*-CY5 probes for 4 h at 37°C, egg chambers were then incubated with primary antibodies overnight at room temperature in 0.3% Triton X-100, 0.2% BSA, 2X SSC. After three washes with 0.05% Triton X-100 in 2X SSC, the egg chambers were incubated with fluorescently labeled secondary antibodies [Alexa Fluor Plus 555, Alexa Fluor 405 (1:1,000; Life Technologies)] for 2 h at room temperature in 0.3% Triton X-100, 0.2% BSA, 2X SSC, and washed and mounted as previously described (Bayer et al, 2015). Nuclear membrane was detected with Wheat germ agglutinin (WGA) CF405S (1:100; Biotium) and actin was detected with Phalloidin DyLight554 (1:200; ThermoFisher). Primary antibodies used were rat anti-Cup (1:1,000) (kind gift from Dr. Allan Spradling, Carnegie Institution for Science), mouse anti-Cup (1:1,000) (kind gift from Dr. Akira Nakamura,Institute of Molecular Embryology and Genetics, Kumamoto University), rabbit anti-Bru1 (1:3,000) (kind gift from Dr. Mary Lilly, National Institutes of Health National Institutes of Child Health & Development) rabbit anti-Osk (1:1000) (kind gift from Dr. Paul Lasko, McGill University) and rabbit anti-GFP (1:1,000) (Millipore).

### Live imaging

Whole ovaries were dissected from well fed females expressing either Bru1-GFP or Cup-YFP in a *wild type* or knocked down (RNAi) background, and individual egg chambers were separated in Halocarbon oil 700 (Sigma-Aldrich) on a #1.5 cover glass. Egg chambers were microinjected with a solution containing a cocktail of five *osk*-specific molecular beacons at a concentration of 200 ng/µl of each probe. Molecular beacons were designed, synthesized, labeled and purified as we previously described (Supplemental_Methods_S2) (Bayer et al, 2018; Bratu et al, 2003). Imaging was initiated within seconds after microinjection, and data acquisition was performed at room temperature on a LeicaDMI-4000B inverted microscope (Leica Microsystems, Buffalo Groove, IL) mounted on a TMC isolation platform (Technical Manufacturing Corporation, Peabody, MA), with a Yokogawa CSU10 spinning disc head and Hamamatsu C9100–13 ImagEM EMCCD camera. The microscopy set up includes diode lasers [491, 561, and 638 nm (Spectra Services, Ontario, NY)], and an Eppendorf Patchman-Femtojet microinjector (Eppendorf, Hauppauge, NY). The images were acquired as 16-bit data files, with 40x/1.25, 63x/1.4 or 100x/1.42 oil objectives (0.385, 0.24 and 0.143 µm/pixel, respectively), using Volocity acquisition software (Quorum Technologies). The second microscope set up was composed of a Yokogawa CSU22 spinning disc head and Hamamatsu ORCA-ER CCD camera, mounted on an inverted Nikon TE2000-U microscope with an 60x/1.49 oil objective. It was equipped with five Obis solid state lasers (405, 488, 514, 561, 640 nm) controlled by an Obis Scientific Remote. Acquisition software: Nikon NIS – Elements v 4.6. The third microscope set up was Leica TCS SP8 WLL. It was equipped with a White Light Laser (470-670nm), a solid-state laser (405 nm), an acousto-optical tunable filter and an 63x/1.4 oil objective. Acquisition software: Leica LAS-X. Optical Z-stacks with a slice thickness ranging from 0.3 to 0.5 µm were acquired using a manual XY-stage with piezo-Z (PerkinElmer), or automated XYZ-piezo stage, respectively. For tracking analysis, 11 to 17 Z-slices of 0.3 to 0.5 µm were acquired every 30 sec for at least 10 min, using the “High-quality” camera mode.

### Image processing, particle detection and tracking

Identical image acquisition, processing parameters and same stage egg chambers were compared with *wt*. Images were deconvolved using AutoQuant X (Media Cybernetics) with adaptive PSF (Blind) using the default recommended expert settings (10 iterations, 10 interval and 20 noise level). Images were processed with ImageJ and Icy softwares (de Chaumont et al, 2012; Rueden et al, 2017). Particle detection, colocalization and tracking analyses for live egg chamber experiments were performed with Icy, using the “Spot detection”, “Colocalization’’ and “Spot Tracking” blocks. For live cell images acquired with the 63x objective, deconvolution and background subtraction were performed, and then spots were detected using the following parameters: scale 2 and 70-110 sensitivity; object-based bright spot detection and size filtering of minimum 4 pixels. Colocalization was determined as a distance of 4 pixels or less from the middle of the 2 particles. Colocalization of *osk* mRNA with either Bru1-GFP or Cup-YFP was measured using an Icy protocol with “Spot detection and Colocalizer’’. Only colocalized particles were used to perform tracking analysis. Tracking parameters were estimated for diffusive and directed motion. Fixed images acquired with the 40x objective were not deconvolved. For fixed images acquired with the 63x or 100x objectives, colocalization analysis was carried out after deconvolution without any further processing using Icy, at scale 1 and sensitivity 40 (*osk* mRNA), 60 (Cup-YFP), 80 (Bru1-GFP), 70 (Me31B-GFP), with minimum size filtering of 3 pixels. All image insets are 2x magnification of the corresponding region indicated with a white rectangle using the Zoom in Images and Stacks macro by Gilles Carpentier available in ImageJ. Pairwise stitching was used in a subset of images in ImageJ (Preibisch et al, 2009). Imaris 9.7 image analysis software surface detection module was used to detect Me31B-GFP signal to calculate the volume and the sphericity of each object detected. Thresholding and sensitivity were carefully determined for each image, objects smaller than 0.15 µm^3^ for early and 0.2 µm^3^ for mid oogenesis were removed from the analysis. Statistical analysis was carried out using Prism 8 software (GraphPad) using unpaired *t* test to calculate the p value, except where otherwise indicated. On the colocalization and mRNA quantification graphs each data point is represented along with mean ± SEM. Normalized fluorescence intensity calculation for neighboring nurse cells: (i - i_min_)/(i_max_-i_min_). Line profiles were performed with a linewidth of 8 µm, 26 µm and 8 µm in Fig. 2G, 3H, and S2C, respectively.

### Western blotting analysis

Ovaries from young females fed on yeast paste for 3-5 days were dissected in Robb’s medium. The samples were lysed with Cytobuster™ Protein Extraction Reagant (Millipore) supplemented with HALT protease inhibitor cocktail. Antibodies used were anti-GFP (1:5,000) (Millipore), anti-Bru1 (1:5,000) (kind gift from Dr. Mary Lilly, National Institutes of Health National Institutes of Child Health & Development) and anti-α Tubulin 12G10 (1:15,000) (DSHB Hybridoma Product 12G10 anti-alpha-tubulin, the monoclonal antibody developed by Frankel, J. / Nelsen, E.M. was obtained from the Developmental Studies Hybridoma Bank, created by the NICHD of the NIH and maintained at The University of Iowa, Department of Biology, Iowa City, IA 52242). HRP-conjugated secondary antibodies were used at 1:5,000 anti-rabbit (Cell Signaling) or anti-mouse (Fisher Scientific).

### RNA isolation and RT-qPCR

Whole ovaries were dissected in Robb’s medium, washed with 1X PBS and kept on ice. As control for the RNAi machinery, we drove RNAi for *mCherry*, a non-endogenous gene in the fly genome. Total RNA was isolated using TRIzol (ThermoFisher) according to the manufacturer’s instructions. RT reactions using 2.25 µg total RNA, Superscript III (Life Technologies) and dT18 oligomers were used to synthesize cDNA. No RT enzyme reactions were used as negative controls. Primers were designed using DRSC FlyPrimerBank for *Drosophila* (Primer sequences: Supplemental_Methods_S3). Primers were acquired from Integrated DNA Technologies. qPCR was performed on white 384-well plates (E&K Scientific, Santa Clara, CA) using Roche Lightcycler 480 (Roche Molecular Systems, Inc.). Each reaction contained 1 µL of cDNA from the RT reaction, 2 µL of primer solution containing 10 µM forward and reverse primers, 5µL SYBR Green I Master mix (5 mL: Roche Diagnostics, Indianapolis, IN) and 2 µL of distilled and deionized H_2_O. Plates were spun down to eliminate air bubbles. Reactions were carried out as follows: 95°C denaturation for 5 min, followed by 37 cycles of 95°C for 20 sec, 58°C for 15 sec. Statistical analyses were carried out using Prism software using unpaired *t* test to calculate the p value.

