## Supplemental Figures and Tables for "Cup is essential for *oskar* mRNA translational repression during early *Drosophila* oogenesis"

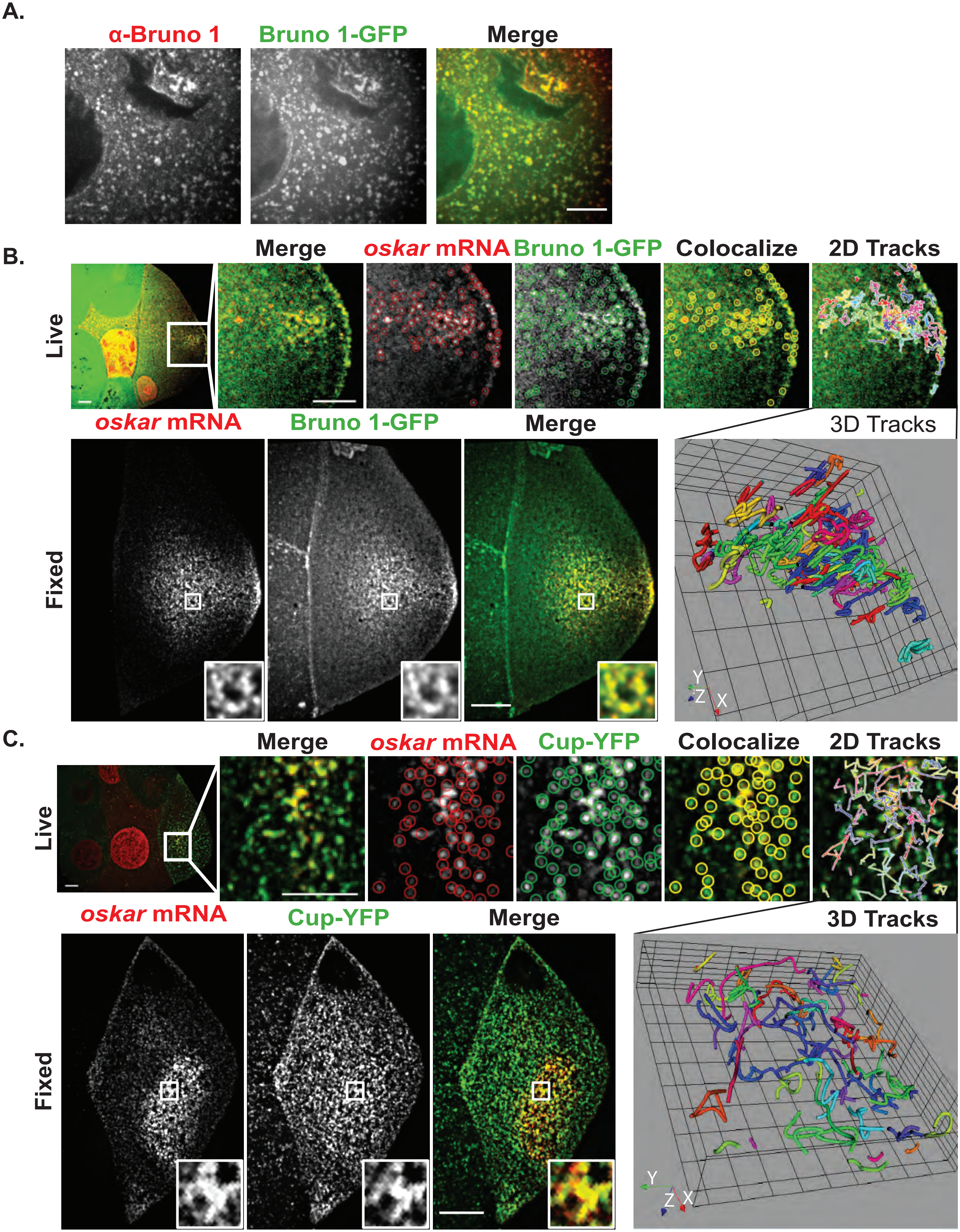
A.
α-Bruno 1
Bruno 1-GFP
Merge
B.
Live
Merge
oskar mRNA
Bruno 1-GFP
Colocalize
2D Tracks
oskar mRNA
Bruno 1-GFP
Merge
3D Tracks
Fixed
Y
Z
X
C.
Live
Merge
oskar mRNA
Cup-YFP
Colocalize
2D Tracks
oskar mRNA
Cup-YFP
Merge
3D Tracks
Fixed
Y
Z
X

Figure S1

**Figure S1. Cup and Bru1 stably associate with *osk* mRNA in the oocyte throughout oogenesis.**

**(A)** Representative image showing colocalization of endogenously tagged heterozygous Bru1-GFP with Bru1 detected via IF in nurse cells of stage 7/8 egg chamber. Image is a single optical slice.

**(B)** Co-visualization of *osk* mRNA and Bru1-GFP in stage 8 live or fixed oocytes. Top panel: colocalization analysis in live egg chambers at the 23 min time point of a 39 min XYZCt series, acquired every 30 sec. Tracks are randomly colored over 15 time points, and are shown as a 2D XYZt-projection (39 min) and a 3D representation. Images are deconvolved XY max-intensity Z-projections of 14 (live) and 16 (fixed) optical slices (0.3 µm each).

**(C)** Co-visualization of *osk* mRNA and Cup-YFP in stage 7 live or fixed oocytes. Top panel: colocalization analysis in live egg chambers at the 7.5 min time point of a 20 min XYZCt series, acquired every 30 sec. Tracks are randomly colored over 10 time points, shown as a 2D XYZt-projection (20 min) and a 3D representation. Images are deconvolved XY max-intensity Z-projections of 14 (live) and 16 (fixed) optical slices (0.3 µm each).

Scale bars, 10 µm.

See also Videos S1, S2, S3, and S4.

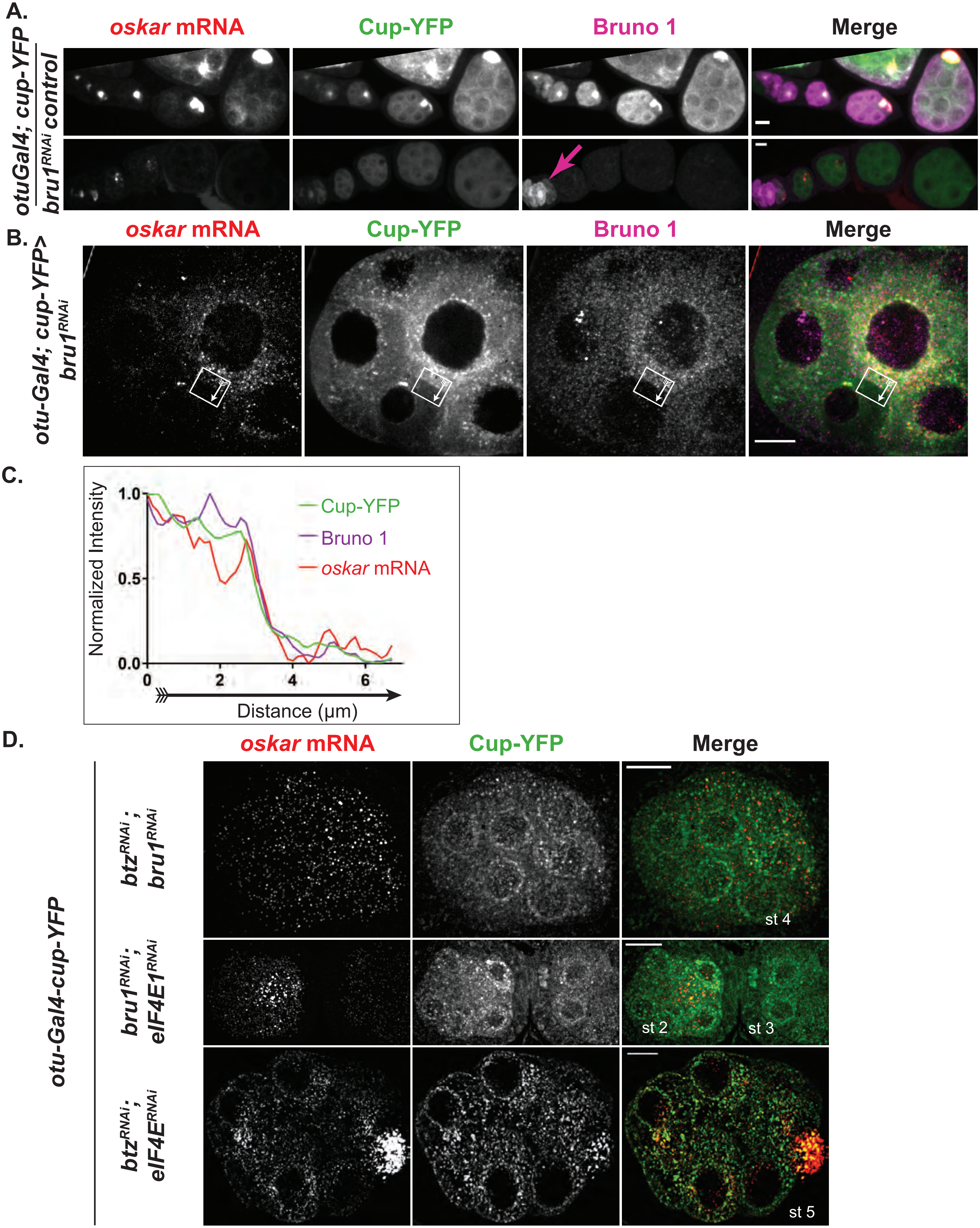
A.
oskar mRNA
Cup-YFP
Bruno 1
Merge
otuGal4; cup-YFP
control
bru1RNAi
B.
oskar mRNA
Cup-YFP
Bruno 1
Merge
otu-Gal4; cup-YFP>
bru1RNAi
C.
Normalized Intensity
1.0
0.5
0.0
0
2
4
6
Distance (μm)
Cup-YFP
Bruno 1
oskar mRNA
D.
oskar mRNA
Cup-YFP
Merge
otu-Gal4-cup-YFP
btzRNAi ; bru1RNAi
st 4
bru1RNAi; eIF4E1RNAi
st 2
st 3
btzRNAi; eIF4ERNAi
st 5

Figure S2

**Figure S2. Cup levels are decreased in *bru1* knockdown egg chambers, and when two other *osk* mRNP members are knocked down, but *osk* mRNA still colocalizes with Cup.**

**(A)** Representative *osk* smFISH and Bru1 IF experiments in egg chambers expressing Cup-YFP in *wt* or *bru1* $^{RNAi}$ backgrounds. Images are XY max-intensity Z-projections of 18 and 11 optical slices (0.3 μm each), respectively.

**(B, C)** Same as Figure 2D including *osk* smFISH experiment.

**(D)** Representative *osk* smFISH experiments in egg chambers expressing Cup-YFP in the indicated RNAi backgrounds (*bru1* and *btz*, *bru1* and *eIF4E*, *btz* and *eIF4E*). Images are deconvolved XY max-intensity Z-projections of 34, 37 and 16 optical slices (0.3 μm each), respectively. Control panel is represented in Fig. 2A.

Scale bars, 10 μm.

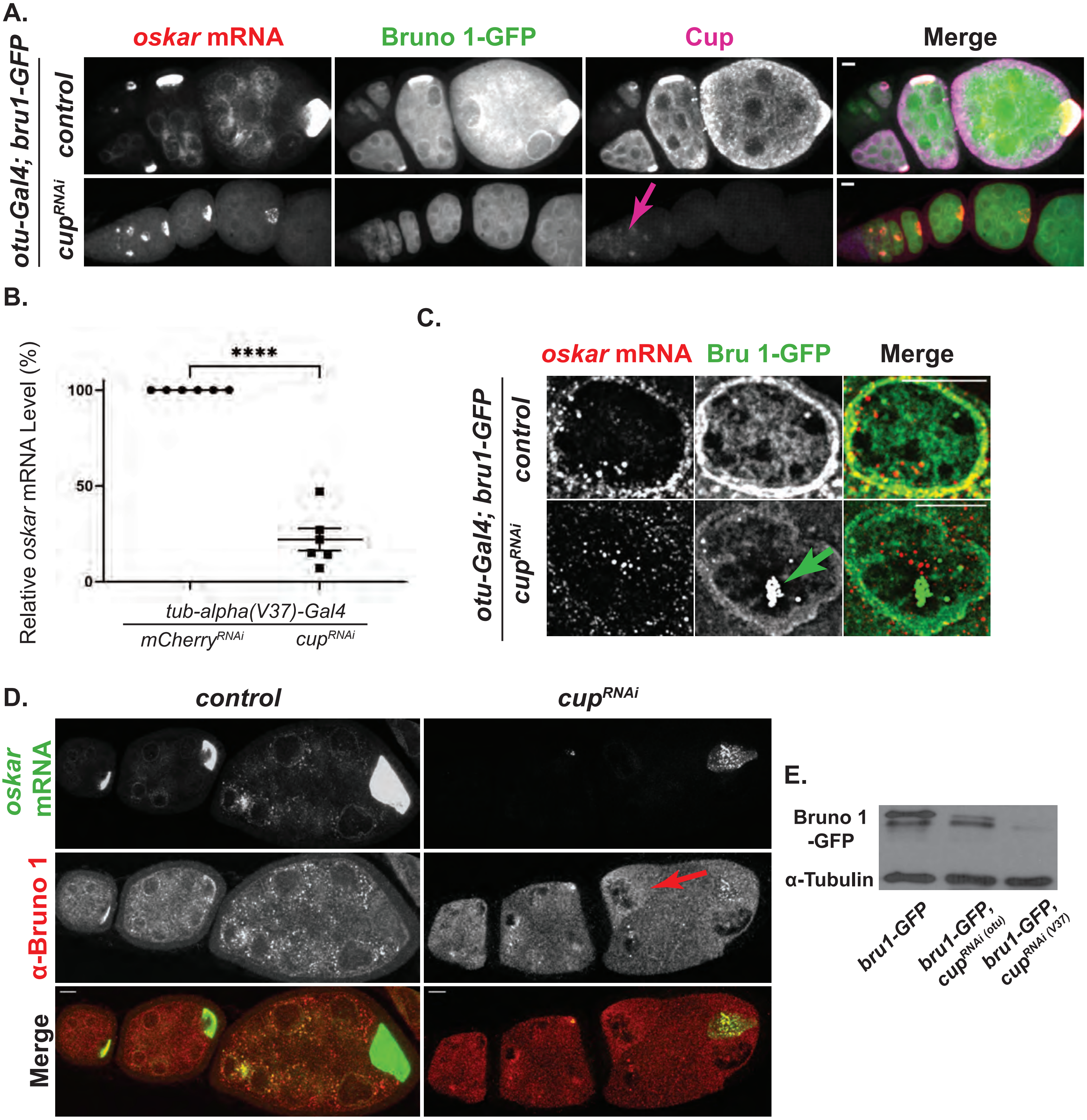
A.
oskar mRNA
Bruno 1-GFP
Cup
Merge
otu-Gal4; bru1-GFP
control
cupRNAi
B.
Relative oskar mRNA Level (%)
****
100
50
0
tub-alpha(V37)-Gal4
mCherryRNAi
cupRNAi
C.
oskar mRNA
Bru 1-GFP
Merge
otu-Gal4; bru1-GFP
control
cupRNAi
D.
control
cupRNAi
oskar mRNA
α-Bruno 1
Merge
E.
Bruno 1 -GFP
α-Tubulin
bru1-GFP
bru1-GFP, cupRNAi (otu)
bru1-GFP, cupRNAi (V37)

Figure S3

**Figure S3. Reduced Cup levels alter Bru1's distribution in the egg chamber, and Bru1 forms large aggregates in the nucleus.**

**(A)** Representative *osk* smFISH and Cup IF experiments in egg chambers expressing Bru1-GFP in *wt* or $cup^{RNAi}$ backgrounds. Images are XY max-intensity Z-projections of 22 and 16 optical slices (0.3 µm each), respectively.

**(B)** RT-qPCR quantification of endogenous *osk* mRNA levels in the indicated backgrounds, normalized to *rp49* mRNA (mean ± SEM ****p<0.0001; $mCherry^{RNAi}$ n=6, $cup^{RNAi}$ n=6).

**(C)** Nurse cell nucleus of *wt* or $cup^{RNAi}$ egg chambers at mid oogenesis. Green arrow highlights the large nuclear aggregation of Bru1-GFP signal in $cup^{RNAi}$ background. Images are deconvolved XY max-intensity Z-projections of 22 (*wt*) and 16 ($cup^{RNAi}$) optical slices (0.3 µm each).

**(D)** Representative *osk* smFISH and Bru1 IF experiments in egg chambers in *wt* or $cup^{RNAi}$ backgrounds. Images are XY max-intensity Z-projections of 10 optical slices (0.3 µm each).

**(E)** Immunoblot against GFP and α-Tubulin using lysates prepared from egg chambers expressing Bru1-GFP in the indicated backgrounds.

Scale bars, 10 µm.

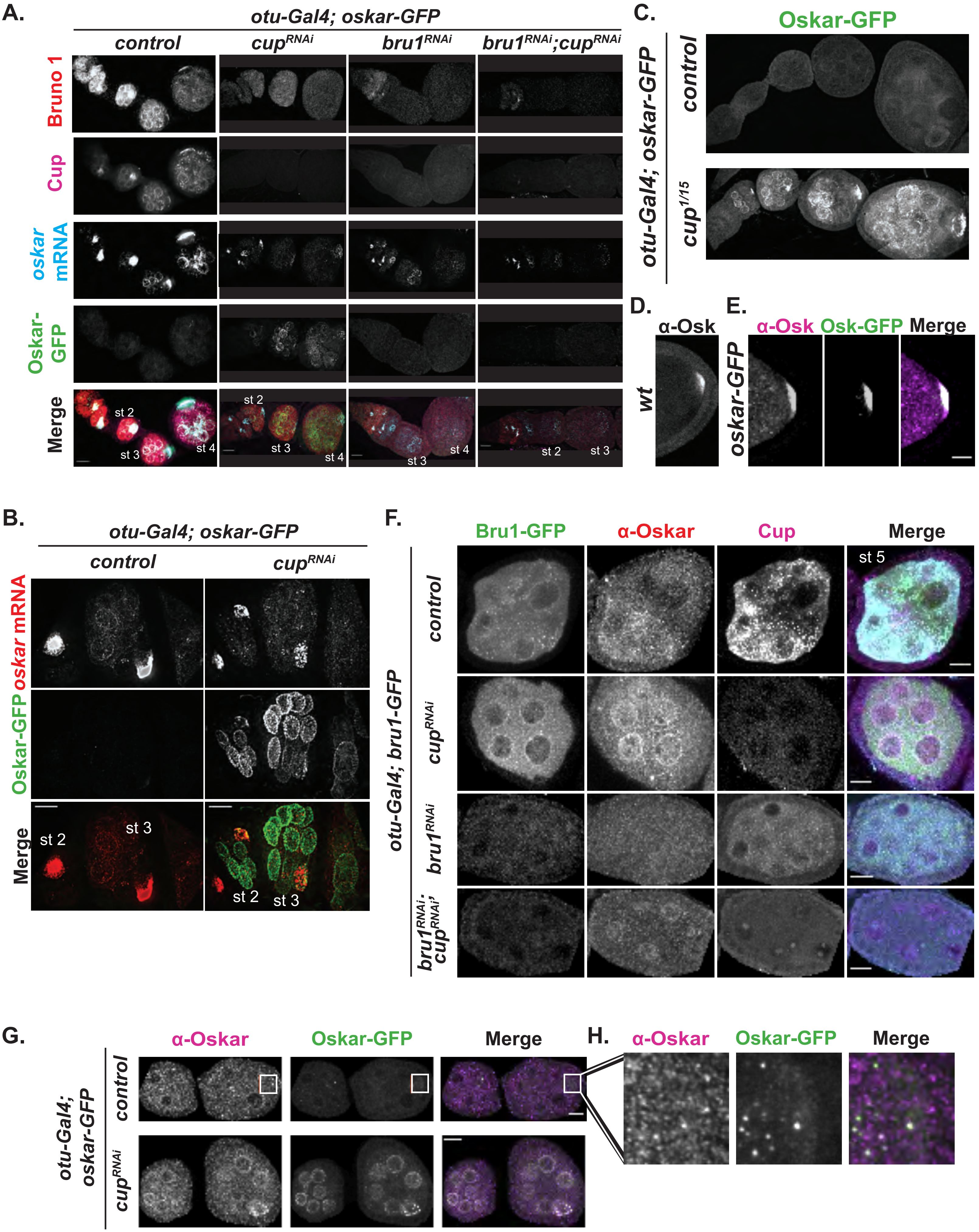
A.
otu-Gal4; oskar-GFP
control
cupRNAi
bru1RNAi
bru1RNAi;cupRNAi
Bruno 1
Cup
oskar mRNA
Oskar-GFP
Merge
st 2
st 3
st 4
B.
otu-Gal4; oskar-GFP
control
cupRNAi
oskar mRNA
Oskar-GFP
Merge
C.
Oskar-GFP
otu-Gal4; oskar-GFP
control
cup1/15
D.
α-Osk
wt
E.
α-Osk
Osk-GFP
Merge
oskar-GFP
F.
Bru1-GFP
α-Oskar
Cup
Merge
st 5
otu-Gal4; bru1-GFP
control
cupRNAi
bru1RNAi
bru1RNAi; cupRNAi
G.
α-Oskar
Oskar-GFP
Merge
otu-Gal4; oskar-GFP
control
cupRNAi
H.
α-Oskar
Oskar-GFP
Merge

Figure S4

**Figure S4. Cup directs translational repression of *osk* mRNA, while Bru1 is involved in its translational activation during early stages of oogenesis.**

**(A)** Representative images of *osk* smFISH and simultaneous detection of Cup and Bru1 via IF in egg chambers expressing Osk-GFP in indicated backgrounds. Images are deconvolved XY max-intensity Z-projections of 94 (*wt*), 85 (*cup*$^{RNAi}$), 77 (*bru1*$^{RNAi}$) and 53 (*bru1*$^{RNAi}$*;cup*$^{RNAi}$) optical slices (0.3 µm each).

**(B)** Representative images of *osk* smFISH experiments in egg chambers expressing *osk-GFP* in indicated backgrounds, showing the distribution of *osk* mRNA and Osk-GFP protein.

**(C)** Representative images of a chain of egg chambers (germarium through stage 5) expressing Osk-GFP in the indicated backgrounds. Images are deconvolved XY max-intensity Z-projections of 20 (*wt*), 22 (*cup*$^{1/15}$).

**(D)** Representative image showing endogenously Osk detected via IF in a stage 9 oocyte. Image is a XY max-intensity Z-projection of 10 optical slices (0.3 µm each).

**(E)** Representative image showing colocalization of endogenously tagged heterozygous Osk-GFP with Osk detected via IF in a stage 9 oocyte. Image is a XY max-intensity Z-projection of 21 optical slices (0.3 µm each).

**(F)** Representative images taken of simultaneous detection of Cup and Osk via IF experiments in egg chambers expressing Bru1-GFP in indicated backgrounds. Images are XY max-intensity Z-projections of 15 (*wt*), 11 (*cup*$^{RNAi}$), 15 (*bru1*$^{RNA}$*i*) and 15 (*bru1*$^{RNAi;}$*cup*$^{RNAi}$) optical slices (0.3 µm each).

**(G)** Representative image showing colocalization of endogenously tagged heterozygous Osk-GFP with Osk detected via IF in the indicated backgrounds of stage 2/3 egg chambers. Images are XY max-intensity Z-projections of 10 (*wt*) and 10 (*cup* $^{RNAi}$).

**(H)** Inset from (G) showing premature Osk-GFP and Osk colocalization in the oocyte of *wt* egg chamber.

Scale bars, 10 μm.

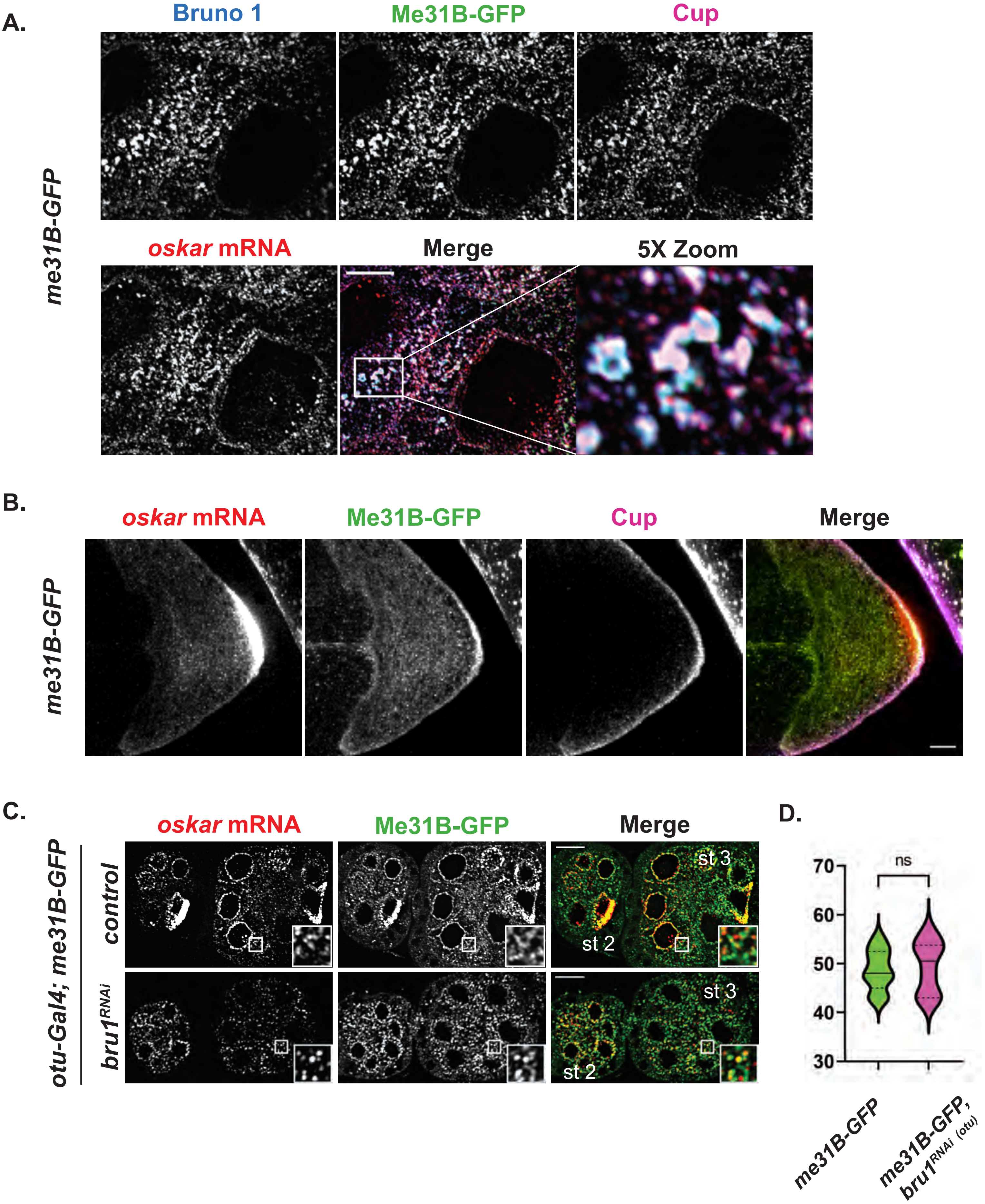
A.
Bruno 1
Me31B-GFP
Cup
me31B-GFP
oskar mRNA
Merge
5X Zoom
B.
oskar mRNA
Me31B-GFP
Cup
Merge
me31B-GFP
C.
oskar mRNA
Me31B-GFP
Merge
otu-Gal4; me31B-GFP
control
bru1RNAi
st 3
st 2
st 3
st 2
D.
ns
70
60
50
40
30
me31B-GFP
me31B-GFP, bru1RNAi (otu)

Figure S5

**Figure S5. Colocalization of Me31B with Cup, Bru1 and *osk* mRNA**

**(A)** Representative images of *osk* smFISH, Cup and Bru1 IF in the nurse cells of a stage 6/7 egg chamber expressing Me31B-GFP. Image is an XY max-intensity Z-projection of 5 optical slices (0.3 µm each).

**(B)** Representative images of *osk* smFISH and Cup IF in the oocyte of a stage 8 egg chamber expressing Me31B-GFP. Image is an XY max-intensity Z-projection of 5 optical slices (0.3 µm each).

**(C)** Representative images of *osk* smFISH in early stage egg chambers expressing Me31B-GFP in *wt* or $bru1^{RNAi}$ background.

**(D)** Colocalization percentage of *osk* mRNA with Me31B-GFP was performed in the indicated backgrounds (*wt* (n=9), $bru1^{RNAi}$ (n=6)).

Scale bars, 10 µm.

**Supplemental_Movie_S1.** Live co-visualization of *osk* mRNA and Bru1-GFP in the nurse cells.

**Supplemental_Movie_S2.** Live co-visualization of *osk* mRNA and Bru1-GFP in the oocyte.

**Supplemental_Movie_S3.** Live co-visualization of *osk* mRNA and Cup-YFP in the nurse cells.

**Supplemental_Movie_S4.** Live co-visualization of *osk* mRNA and Cup-YFP in the oocyte.

**Supplemental_Movie_S5.** Live co-visualization of *osk* mRNA and Bru1-GFP in $cup^{RNAi}$ egg chambers, for a 38 min XYZCt series. Red and green circles indicate *osk* mRNA and Bru1-GFP, respectively. Images are XY max intensity projections of 7 optical slices (0.3 μm each). Scale bars represent 10 μm.

**Supplemental_Table_S1.** Probe sequences used in FISH experiments.

**Supplemental_Table_S2.** Molecular Beacon probe sequences used in live cell imaging.

**Table S1. Probe sequences used in smFISH experiments for *oskar* mRNA.**
.

| Sequence |
|---|
| ttgctctcgatgatggtcat |
| ttcctcgcgcacggatatat |
| cctcactatctatatcggga |
| tatcgtgattccattctggg |
| gatattcactcttgatgctc |
| aatggattgcccgtcagttt |
| aaatccgtcacgttgtcgtg |
| cacattgggaatggtcagca |
| ttgaagatccgcttaccgga |
| gttcttcaggctcgctttca |
| ttctggttgagcaccatatc |
| tgctcgtggtatgttctcca |
| tgaagggattcttccagtag |
| ttcatccgttatcttgggca |
| tcattggccatttgatgcag |
| ttatcctggtagcaccagtt |
| ggcttgctggtagaaattgt |
| gcatttttggcgcatttacg |
| ctgtagatgttgatgggtac |
| aattgattggttcctctggg |
| ttttctggctttgggttctg |
| atggcggttttcagtcggtt |
| aatggatgcacaaagatgcc |
| ggcgtctcttcattatgttc |
| ttaaaatcgttggcgtgggc |
| gaatcgttgtaggttccact |
| atccgagttaatcgtcagca |
| aagtccagcagataggcatc |
| aaaatcatcgcccataagcg |
| tccattcgggcgagatatag |
| gacgcctaaatcggcatttt |
| tacacaaagtcctgactgca |
| aaacgatttcgggcgccatt |
| ctcgggtaattggactcttt |
| ccgatattgacgatgatctg |
| aaaggcttgccgcgcataat |
| gaaaatcgtgctcgatctga |
| tgcatctccttgatcagtag |
| aatcggcaccaatcgcatat |
| atcgtgacaatagttgccca |
| gaatcggtagattttgtcac |
| gcattcgcttcggataaact |
| tgtcgatgacctttaggtga |
| tcgttgattagacaggagtg |
| aagcaatcaaatcgcaccac |
| ttccttggaaccggtgactt |
| ccgattttgttccagaacag |
| aataacttgcagtacgcgct |

**Table S2. Molecular Beacon probe sequences used in live cell imaging to detect *oskar* mRNA.**

**Sequences**

5'- cgaccGACUUAAGAUAAUAGGUUUUGGCGggucg -3'
5'- GGACC GAGUGGACGAGAAGAGUCUGG GGUCC -3'
5'- CCUGG UCGCUGGUGCGCUCUUUCUGGUU CCAGG -3'
5'- CCAGC UUGCUGGUAGAAAUUGUUGAGAU GCUGG -3'
5'- gcugcAAAAGCGGAAAAGUUUGAAGAGAAgcagc -3'
